# Right on time: prediction-errors bidirectionally bias time perception

**DOI:** 10.1101/766204

**Authors:** Ido Toren, Kristoffer Aberg, Rony Paz

## Abstract

The brain updates internal representation of the environment by using the mismatch between the predicted state/outcome and the actual one, termed prediction-error. In parallel, time perception in the sub-second range is crucial for many behaviors such as movement, learning, memory, attention and speech. Both time-perception and prediction-errors are essential for everyday life function of an organism, and interestingly, the striatum was shown to be independently involved in both functions. We therefore hypothesized that the putative shared circuitry might induce behavioral interaction, namely that prediction-errors might bias time perception. To examine this, participants performed a time-duration discrimination task in the presence of positive and negative prediction-errors that were irrelevant and independent of the main task. We find that positive/negative prediction-errors induce a bias in time perception by increasing/decreasing the perceived time, respectively. Using functional imaging, we identify an interaction in Putamen activity between encoding of prediction-error and performance in the discrimination task. A model that accounts for the behavioral and physiological observations confirms that the interaction in regional activations for prediction-errors and time-estimation underlies the observed bias. Our results demonstrate that these two presumably independent roles of the striatum can actually interfere or aid one another in specific scenarios.

---

Time perception in the sub-second range is essential for human behaviors such as speech, movement, attention, memory and other functions (1–5). The subjective perception of time is affected by physical attributes of the stimulus such as spatial frequency, statistics, target type, saliency and intensity (6–9), but also by motivational states and emotional content (10). Mainly, studies have observed that the duration of a stimulus with an emotional content, such as a face with an emotional expression or an arousing image, is perceived as longer compared to other neutral stimuli (11–14).

Accurate and efficient processing of emotions, reinforcements, and punishments, is critical for survival and mediates associative learning. The acquisition of action-outcome associations is driven by the mismatch between the predicted and the actually received outcome, termed prediction-error (PE) (15–18). Outcomes that are better than expected, termed here positive prediction-errors (PE+), or worse than expected, termed negative prediction-errors (PE-), respectively increase or decrease the propensity to perform the associated action.

Time perception and associative learning were classically considered to be largely independent processes, yet recent ideas suggest they might be related (19–22). Time perception engages striatal brain regions that receive dopaminergic projections from the midbrain (23–27), and neural correlates of PEs have been found in dopaminergic midbrain neurons and their striatal projection sites (28–31). Moreover, PE+ and PE− have been associated with increased and decreased activation of dopaminergic neurons, respectively (15), confirmed by similar findings in human studies (32, 33), and a recent study found that the perceived duration of a stimulus could be increased/decreased by activation/deactivation of dopaminergic neurons (34). Not surprisingly, time-perception is compromised in Parkinson-disease and other basal-ganglia related disorders(35–38).

The finding that striatal and dopaminergic circuits play a major role in both functions: time-perception and coding of prediction-errors, suggest that the two might interact and could either interfere or aid one another. We therefore hypothesized that signed prediction errors would decrease or increase the perceived duration of a stimulus, and that such bias would be associated with differential striatal activity.

To examine this, participants determined which of two sequentially presented images is of the longer duration in a 2-alternative-forced-choice paradigm (2AFC, Fig. 1A). There were two types of trials: ‘Short-Long’ (SL, 50% of all trials) where the duration of the first image was shorter, and ‘Long-Short’ (LS, 50%). The difference in time duration between the two images varied across trials (Δt, 0-26.6% corresponding to 0-133ms). To impose an independent PE and test its impact on time perception, each image was overlaid by a number that determined the monetary gain or loss. The first image was lways presented with a zero (0), whereas the number on the second image could be negative (PE−; 20% of trials), positive (PE+; 20%), or 0 (PE0; 60%). The numbers comprised of values from −5 to +5 in gaps of 0.5 and were pseudorandomly drawn from a homogenous distribution. The outcome of a trial was determined only by the number presented on the second image and was independent of the discrimination performance (correct/incorrect). Two studies were conducted using the same procedure. A first behavioral study established the behavioral bias that results from the influence of PEs on time perception (n = 18), and a consecutive fMRI study was carried to replicate behavior and elucidate the neural correlates (n = 35).

**Figure 1.**
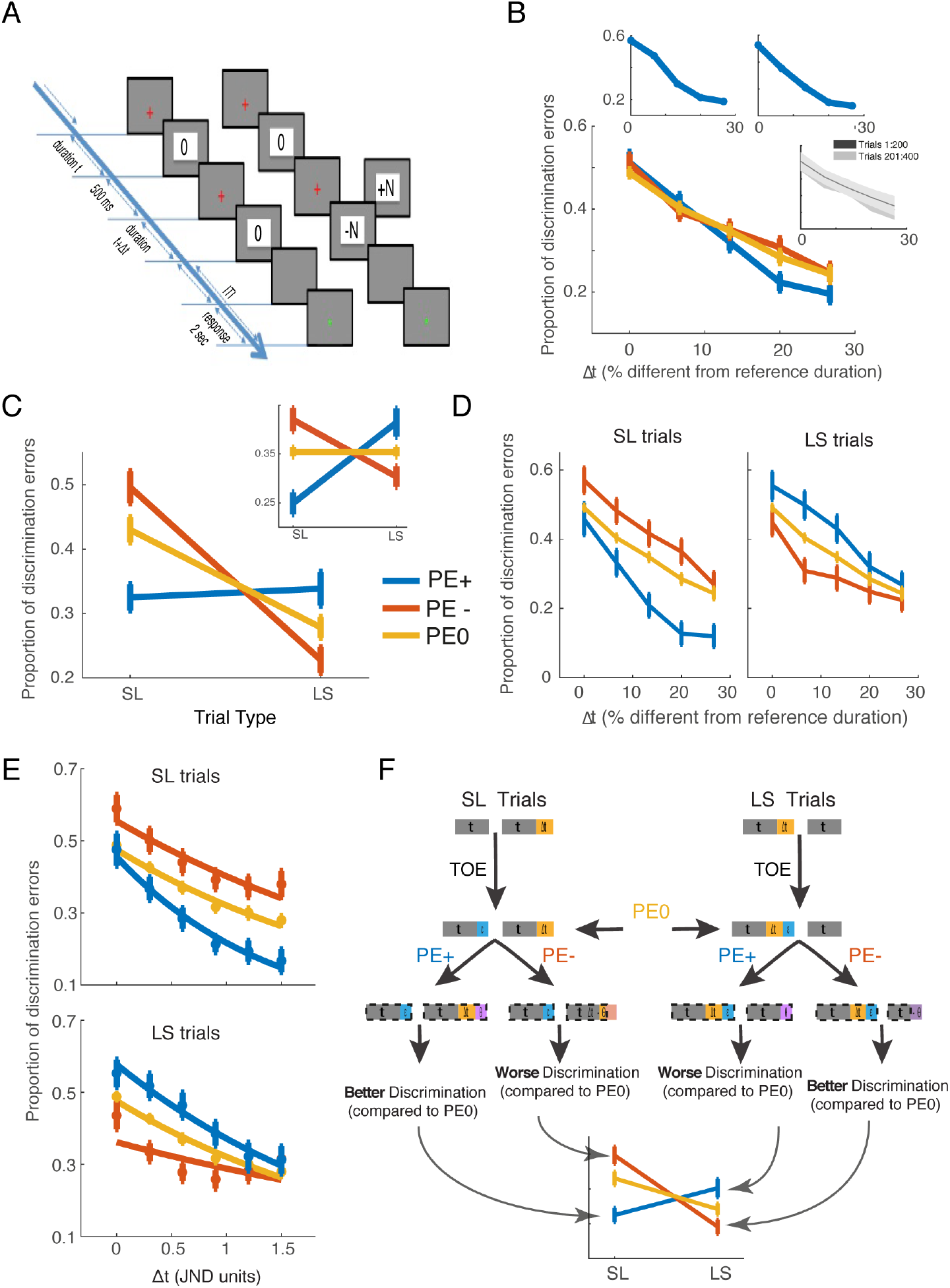
Prediction-errors bidirectionally bias time perception. (A) The 2AFC time-discrimination task. Example of Short-Long (SL) trial with PE0 (left) or with PE+ / PE− (right). (B) Proportion of discrimination errors as a function of objective time difference between the two images (Δt), averaged across all trials separately for each PE type. Insets show two individual subjects (left), and the proportion of discrimination errors in first and second half of the experiment (right, averaged over all subjects), indicating no change in perception throughout the experiment. (C) Proportion of discrimination errors as a function of PE type and trial type. An interaction between PE-type and trial-type is evident as PE+ and PE− bias performance in opposite directions relative to PE0 trials. Inset: same after correction to individual Time-Order Error (TOE). (D) Proportion of discrimination errors as a function of time difference between images (Δt), presented separately for Long-short and Short-long trials and corrected for Time-Order Error (TOE). PE+ and PE− bias performance in opposite directions relative to PE0, for all values of Δt. (E) Proportion of discrimination errors as a function of subjective JND, corrected for TOE. Data fitted to a logistic function after JND normalization. (F) Schematic representation of the bias effect and the model. Right and left sides of the scheme represent Long-short and Short-long trials, respectively. Rectangles represent perceived duration of the images, with colors indicating different factors affecting the perceived duration: Gray rectangles represent the objective reference duration (t); Yellow denotes the time difference Δt added to the first or second image of LS or SL trials, respectively; Blue represent the change in the first image due to the TOE; and finally, Pink represent the bias due to PE (*θ*), added or subtracted for PE+ or PE−. The last row of the scheme therefore shows the prediction for the perceived duration, denoted by a dashed rectangle, leading to the observed behavioral results.

## Prediction-errors interact with time discrimination

We first grouped outcomes into one positive and one negative group, and a three-way repeated measures ANOVA was conducted with factors Trial-type (LS, SL), PE-type (PE−, PE+, PE0), and Δt (0-26.6% corresponding to 0-133ms). The dependent variable was performance measured as the proportion of correct duration discriminations in each condition. Because discrimination is easier for larger values of Δt, we indeed found a significant main effect of Δt (Fig. 1B, *F*_4,208_ = 98.7, p < 10^−46^). Reassuringly, there was no significant interaction between PE-type and Δt (*F*_8,416_ = 1.9, p = 0.06), nor was the three-way interaction (*F*_8,416_ = 125, p = 0.25).

Yet, in line with our hypothesis, we found a strong and clear interaction between PE and Trial-type (Fig. 1C; *F*_2,104_ = 19.04, p < 10^−7^). This main result was robust for each group separately (behavior-only and fMRI), and also when dividing the PE into different magnitudes (Figs.S1,S2,S3; see Supp.Info. for detailed statistics).

In sum, we found that prediction error interacts with time perception to influence discrimination judgments.

## An opposite effect of PE+ and PE− on perceived duration

We next examined what drives this interaction. In our design the PE occurs only when the second stimulus appears. Therefore, in SL trials when the value induces a PE+ or a PE− (PE+/PE−), if the image is perceived as longer/shorter, it would lead to a perceived larger/smaller Δt between the two stimuli resulting in easier/harder discrimination and hence better/worse performance. For LS trials, PE+/PE− would induce the opposite change in perceived Δt thus resulting in an opposite effect on performance.

Accordingly, we observed better performance in SL trials for PE+ vs. PE− (proportion of discrimination errors; mean PE+ = 0.32, mean PE− = 0.49, post-hoc Tukey, p < 10^−5^) and compared to PE0 (mean PE0 = 0.43, post-hoc Tukey, p = 0.0001). Correspondingly, trials with PE− showed worse performance compared to PE0 (post-hoc Tukey, p = 0.02). The exact opposite was observed in LS trials: performance was worse in PE+ compared to PE− trials (mean PE+ = 0.33, mean PE− = 0.22, post-hoc Tukey, p = 0.0004) and marginally compared to PE0 trials (mean PE0 = 0.275, post-hoc Tukey, p = 0.06). Correspondingly, trials with PE+ showed better performance compared to PE0 (post-hoc Tukey, p = 0.02). Together, these results suggest that PE+ and PE− induce an increase and decrease in the perceived stimulus duration, respectively.

## The contribution of order effects and individual thresholds

Previous studies have identified that a second consecutive stimulus may bias the perceived duration of the first stimulus in a discrimination paradigm, termed Time-Order-Effect (TOE) (39, 40). In our paradigm the TOE would induce more errors in SL trials because the first image would be perceived as longer, and the effect should diminish as Δt increases (‘easier’ trials). This was replicated in our subjects as evident by a main effect of Trial-type (Fig.1C; Fig.S5; *F*_1,52_ = 26.19, p < 10^−5^, due to better performance in LS vs. SL); and an interaction between Trial type and Δt (Fig. S4A, *F*_4,208_ = 11.3, p < 10^−7^), due to better discrimination in harder vs. easier trials (i.e. small vs. large Δt) in LS compared to SL trials (Fig.S5, mean difference = 0.2 in harder and 0.08 in easier trials).

To account for this, we quantified the individual TOE as a function of Δt (methods) and normalized all performance measures accordingly (Fig.1C-Inset; Fig.1D; Fig.S5). As expected, the main effect of Trial Type was no longer significant (*F*_1,52_ = 0.5, p = 0.48) nor was the interaction between Trial type and Δt (*F*_4,208_ = 0.75, p = 0.55), whereas all other results, and importantly the significant interaction between PE and Trial-type, did not change (Fig.1D,C-inset, PE x Trial-type: *F*_2,104_ = 19.04, p < 10^−7^).

Next, we accounted for potential individual differences in perceptual thresholds. To do so, we normalized the objective Δt by each subject’s just-noticeable-different (JND), allowing a comparison across subjects. The JND was measured independently of the main paradigm (and confirmed using several approaches, Fig.S6B, methods). All aforementioned findings were replicated using the normalized psychometric curves, and importantly the significant interaction between PE and Trial-type (Fig.1E, PE x Trial-type: *F*_2,96_ = 17.26, p < 10^−6^; Fig. S1E,S2E,S3E).

In addition, we validated that participants had similar JND thresholds for PE0, PE+, PE− conditions, and also before, during and after the main experiment (Fig. S6A). Importantly, we tested that value alone, i.e. without a PE yet similar monetary outcomes, does not induce a similar bias (Fig. S6A). This indicates that the bias is not a result of outcome-value, adaptation or learning processes, and strongly suggests that PE is indeed the source.

## Modelling the PE-time bias

To provide further insight to how prediction-errors bias time-perception on a trial-by-trial basis, we developed a model to account for the observed findings. The model considers the main factors that affect the perceived duration: the objective time difference between images (Δt), the bias due to PE (denoted as θ), and the TOE (denoted by ε). According to the main findings, PE− decreases the perceived duration of the second image, which decreases/increases the perceived difference in SL/LS trials, respectively; and an opposite bias occurs in PE+ trials (Fig. 1F).

In the model, the expected value (EV) is estimated and updated on a trial-by-trial basis (Fig. S4B) using the transient prediction-error (i.e. the difference between the expected and the actual value) and modulated by a learning rate. The probability of making a correct duration discrimination is then modeled by a logistic function (4, 41, 42) and fitted individually for each subject. We found that the best fit was obtained with a model that includes the above-mentioned parameters (Fig. 2A, Fig. S2F, Fig. S3F; see methods for parameter selection). Validity was further confirmed by observations of the individual model-derived error-probability vs. the actual performance (Fig.2B-left) and the model-derived value vs. the actual outcome (Fig. 2B-right), as well as a tight correlation between the model-derived individual PE+/PE− in correct vs. incorrect trials (fig.2C, Pearson correlation, r = 0.95, p < 10^−28^), and between the model-derived TOE (ε) and the actual TOE measured directly and independently on PE0 trials (Fig. 2D; Pearson correlation; behavior-only group: r = 0.73, p < 10^−3^; fMRI group: r = 0.83, p < 10^−5^).

**Figure 2.**
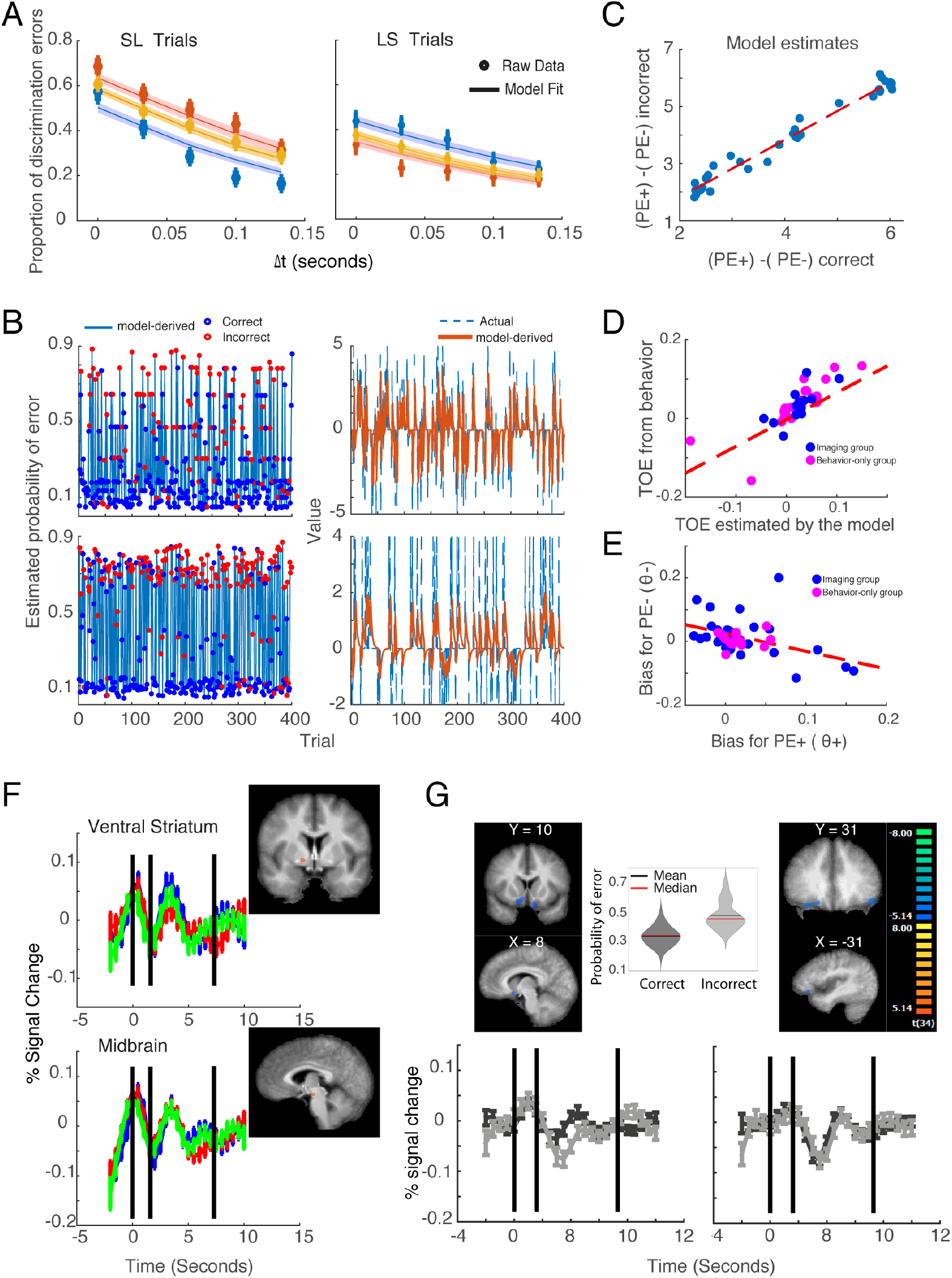
Modelling the PE-time bias. (A) Model fit to individual behavioral data, averaged over all subjects. (B) Two subjects’ (rows) behavior and model fit. Left: model-derived probability of error, with blue/red circles indicating actual correct/incorrect discrimination, respectively. Right: actual outcome overlaid on model-derived expected value. (C) Difference between model-derived PE values is correlated between correct and incorrect trials, validating individual model-derived values. (D) The correlation between TOE from the model fit and TOE computed directly from behavior (quantified as the intersection between performance in PE0 trials and 0.5 proportion of discrimination errors at Δt = 0). (E) The correlation between the bias in time discrimination as a result of PE+ and PE− trials, estimated separately. The bias was not significantly different between PE+ and PE− trials (observed here in the correlation, see text for additional statistics). (F) ROIs in which activation correlates with trial-to-trial model-derived prediction-error, in PE+ trials, displayed on an average brain. Time courses represent mean % signal change extracted from the ROIs ± SEM. Black vertical lines represent trial onset, trial offset and average onset of next trial, respectively. Activation was set to statistical threshold of q < 0.055 for visualization. (G) Activations in ventro-lateral PFC / OFC (right) and NAcc/ventral-ACC (left) was found to correlate with model-derived probability for correct discrimination. Average activation for correct and incorrect choices is plotted next to the ROI, demonstrating higher activation for correct discriminations. Inset shows the mean probability for discrimination error across participants, which is significantly higher for the actual incorrect judgments.

The bias values calculated for PE− and PE+ (θ(PE+), θ(PE−)) were very similar in magnitude and different mainly in sign (Fig. 2E; Pearson correlation; r = 0.65, p < 10^−6^; Paired t-tests comparing θ(PE−) to θ(PE+): all subjects, *t*_53_ = 0.15, p = 0.87; behavior-only group, *t*_17_ = −0.01, p = 0.99; fMRI group: *t*_34_ = 0.16, p = 0.87). Importantly, the average value of the bias (θ) was not different when estimated separately for the behavior-only group and for the fMRI-group (two-samples t-test; *t*_51_ = −1.1, p = 0.26). Finally, to further validate the model accuracy, notice it estimates a PE per trial that is continuous and can take any value. We therefore arbitrarily defined PE0 as trials with |PE|<0.1, and PE+ or PE− for all other positive/negative values, and re-computed the statistics as aforementioned for the behavioral data. This replicated the main finding of interaction between trial-type and PE (as well as other findings, Supp.Info), and was robust to different choices of threshold ranging from 0.005 to 0.5, when 0.2 provides roughly the same proportion of PE0 trials as in the experimental design.

## Brain activity tracks trial-by-trial PE and performance

We first validated that using the model-derived trial-by-trial PE as parametric modulators indeed identify the classical PE network, including the ventral striatum, Midbrain, and dorsal-ACC activations during PE+ trials (Fig.2E, Fig.S7, Table.S2).

Then, we note that the probability for a discrimination-error is estimated by the model on a trial-by-trial basis, and could therefore reflect fluctuations of uncertainty in the perceptual task. We used the estimated probability for each subject and in every trial and found a bilateral negative correlation, namely higher activation when error-probability is low, in the PFC Broadman area 47 (Fig. 2F), a region implicated in state representation (43), confidence (44), and value (45). The same activity pattern of negative correlation with error probability was observed in the border between the Nucleus Accumbens (NAcc) and ventral-ACC (Fig. 2F), a region that was shown to track confidence of prediction errors in a perceptual decision-making task (46). A positive correlation, namely higher activation for higher error-probability, was found in the caudate tail and Right Inferior frontal gyrus, regions previously shown to track trial-by-trial prediction of observers’ categorical perceptual decisions (47) (Table S3). These results further strengthen the model validity and show it integrates prediction-error and time estimations in our task to capture variability in perceptual judgements.

## Putamen activity corresponds to the PE-time bias

Because both time-perception and prediction-error were shown to involve striatal activity and likely depend on dopaminergic modulation, we hypothesized that interaction in the striatum would underlie the bias identified here. To test this, we conducted a whole-brain analysis using two-way ANOVA with factors PE-type (PE+, PE−, PE0, model-derived), and performance (incorrect/correct); all results were corrected for multiple comparisons by non-parametric permutation tests for maximum cluster size (48, 49). This therefore looks for regions that potentially represent the effect of PE leading to difference in performance (correct/incorrect), namely the bias.

We found a significant interaction between PE-type and performance in the Putamen (Fig.3A). During PE+ trials, activation in the Putamen was increased for incorrect compared to correct discriminations, whereas during PE− trials the Putamen was de-activated for incorrect compared to correct judgements. To further confirm that the interaction was driven by PE+ and PE−, we repeated the analysis but without including PE0 trials and found a similar interaction in the Putamen.

**Figure 3.**
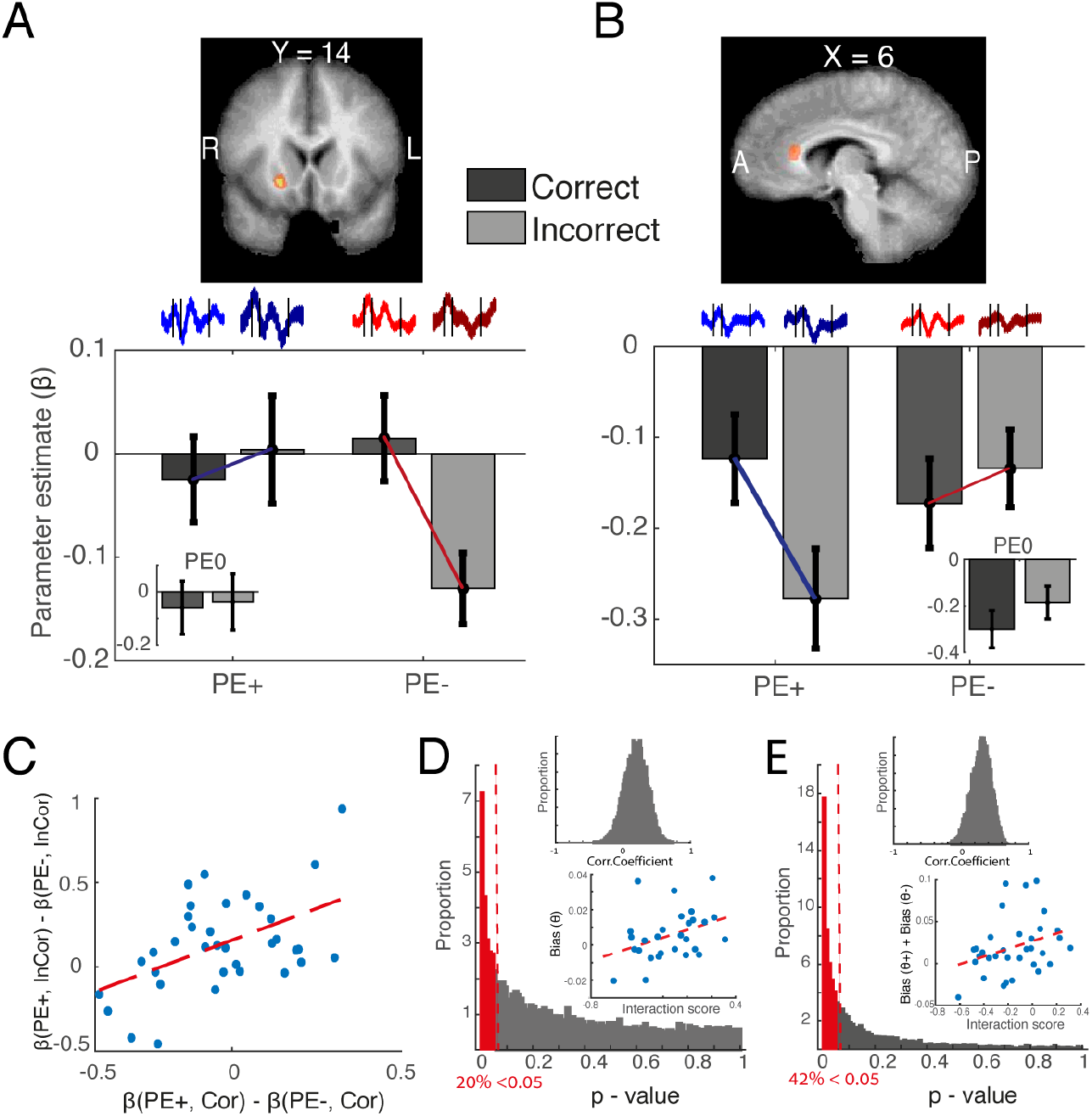
Putamen activity underlies PE-time interaction and behavioral bias. (A) Activation in anterior right putamen is associated with the interaction between PE type (PE+, PE−, PE0) and time discrimination accuracy (correct or incorrect responses). ROIs presented on an average brain, and the mean parameter estimates (beta) for PE type and performance, extracted from these ROIs. Also shown are the corresponding time-course activations. Error bars represent SEM. (B) Same as (A) for the dorsal anterior-cingulate-cortex (dACC). (C) Individual subject Putamen activations (extracted from the above ROIs) for PE+/PE− in incorrect vs. correct discrimination trials, indicating an interaction between PE type and performance at an individual level. (D) Individual interaction-score of Putamen activity is correlated with the individual behavioral bias (θ). Shown is the distribution of p-values for the correlation between the behavioral bias and the individual score of interaction in Putamen activity (20% had a p<0.05, dashed red line and red bars). Distribution was obtained by bootstrap of resampling with replacement from the 35 subjects and correlating individual bias (θ) with the individual activation interaction score in every itertaion. Upper-inset show the distribution of correlation coefficients. Lower-inset shows the correlation for the original data (outliers removed; note the bootstrap to obtain the distribution was done w/o removing any outliers). (E) Same as in (D) but when the model includes a separate bias for PE+ and PE− (θ+ and θ−). Although the two were correlated hence suggesting that a bias is a personal trait that differs between PE+ and PE− mainly in sign (Fig.2E), using the two revealed an even closer match between behavioral bias and the activation pattern in the Putamen (42% had a p<0.05, dashed red line and red bars).

A significant interaction was also found in the dorsal-ACC (Fig.3B, Table.1). Moreover, the individual difference in Putamen activity between PE+ and PE− was correlated across correct and incorrect trials (Fig.3C, r = 0.45, p = 0.005), indicating that Putamen activity tracks these factors at an individual level as well. A similar individual relationship was found in the dACC when considering separately PE+ vs. PE0 and PE0 vs. PE− (Fig. S8; PE+/0: r = 0.37, p=0.03; PE−/0: r = 0.6, p<0.0001).

**Table 1.** Significant ROIs obtained from the interaction between PE-type and time discrimination performance.

| **Region (BA)** | **F-value**<br>**(2,68)** | **x y z**<br>**(Talairach)** | **Cluster size** |
| --- | --- | --- | --- |
| *<u>PE (PE+, PE-, PE0) x</u>*<br>*<u>Correctness</u>* |  |  |  |
| **Right Anterior Putamen** | 14.7 | 21 14 1 | 239 |
| **dorsal ACC (32)** | 14.8 | 6 26 16 | 491 |
| Left Inferior Frontal Gyrus (9) | 8.36 | -33 11 28 | 403 |
| Left Middle Frontal Gyrus (9) | 7.52 | -42 14 40 | 221 |
| Left Middle Temporal Gyrus<br>(21) | 11.31 | -45 -13 -11 | 531 |
| *<u>PE (PE+, PE) x Correctness</u>* |  |  |  |
| Right posterior Insula | 5.91 | 48 -10 10 | 193 |
| **Right Anterior Putamen** | 8.49 | 21 14 1 | 232 |

Finally, to further establish a link between Putamen activation and the behavioral PE-time bias, and at an individual level, we devised an interaction-score that quantifies the interaction in Putamen activity of PE+/PE− with the performance (correct/incorrect), at an individual level (methods), and correlated this activity score with the individual behavioral bias (Fig.3D,E: lower-insets). To establish robustness of this measure we performed a bootstrap by resampling a different set of subjects every time and re-calculating the correlation between the Putamen interaction-factor and the behavioral bias for each subset of subjects (methods). The distribution of the correlation coefficients and their respective p-values was highly skewed and significant (Fig.3D,E, p<0.001 for all, Fisher’s test). These results confirm and suggest a direct link between interaction of activity in the Putamen for the two factors, PE and duration, and the bias that PE induces on time perception.

## Conclusions

Our findings provide first evidence that prediction-errors bias time-perception, atleast in the sub-second range, and shed light on the neural circuits that underlie the behavioral interaction between these two fundamental computations. At the behavioral level, we found that the mismatch between expected and actual outcome, i.e. the prediction-error, results in mis-estimation of the presentation time and leads to incorrect judgement of duration. Interestingly, we identified a bidirectional effect, where a positive prediction-error - receiving more than expected - results in over-estimation of duration, and negative prediction error - receiving less than expected - results in under-estimation of duration.

These findings re-open discussions surrounding the idea that arousal alone, for example during an emotional stimulus which is either negative (10, 50, 51) or positive (52, 53), dictates perception to be of longer duration. The findings further suggest re-consideration of the idea that predictability alone may cause the duration to be perceived as shorter (54, 55). Alternatively, we find a more complex relationship where a signed prediction-error that is processed in timing-related circuits, cannot be accounted by absolute (attention-like) prediction-error signal. It is possible that an integrative mechanism that combines opposite patterns of attention due to saliency (56), valence (57) and/or amount of information passing (58) together with unpredictability(7, 8), could eventually account for our findings.

At the neural level, it was recently shown that manipulating midbrain dopamine (DA) neurons affects time perception (34). While in-line with these reports, our findings suggest that midbrain neurons might not be the locus of this effect, and the signals are integrated and interact in overlapping circuits downstream in the striatum. This is also in-line with results showing that attention to time correlates with activation in the putamen (59). The trial-by-trial variability in probability of correct discrimination, a measure indicating uncertainty that changes according to task difficulty, was found to correlate with activations in networks underlying state, confidence and uncertainty during perceptual decision-making (45–47, 60, 61), indicating that our experimental design and computational model capture perceptual uncertainty in the brain, but one that comes from prediction-errors and their signed directionality.

Because both these processes, prediction-errors and duration-estimation, are essential for functional and cognitive capabilities, the impact of such biases can be of importance during daily life. Learning and memory formation rely on ongoing computations of predictions-errors as well as on estimating durations. Therefore, neural interactions in the striatum can either improve or harm these processes, and the overlap in activity we observe here and that leads to the behavioral bias can be either an evolutionary benefit or a harmful by-product (e.g. by biasing temporal-difference learning(20)). Specifically, we suggest and hypothesize that maladaptive learning can stem from abnormal neural interactions in striatal circuits during formation of internal representations of both time-duration and learning-related errors. In turn and in extreme cases, these interactions can lead to trauma, anxiety, and other experience-dependent psychopathologies(35, 62–65).

## Acknowledgments

We thank Dr. Edna Furman-Haran and Fanny Attar for MRI procedures. This work was supported by a Joy-Ventures grant, ISF #2352/19 and ERC-2016-CoG #724910 grants to R. Paz.

## Author contribution

I.T. and R. P. designed the study. I. T. performed the experiments and analyzed the data. I. T., K.A. and R. P. wrote the manuscript.

## Materials and Methods

### Experimental design

#### Participants

Eighteen (18) healthy participants (5 males) participated in the behavior-only group and 35 healthy right-handed participants (15 males) participated in the imaging group (fMRI). All participants had normal or corrected-to normal vision and reported no attention deficit hyperactivity disorder (ADHD). Informed consent was obtained from all participants prior to the experiment. One participant in the imaging group was unable to complete the fMRI scan due to unexpected physical influences of the magnetic field and another participant could not complete the scan due to extensive movements, both had been excluded.

#### Visual stimuli

White images with black numbers displayed in the center of the image were presented on a gray background. The images were identical in size, presented in the center of a 21” screen with refresh rate of 60 Hz (lag smaller than 1ms), and spanning a visual angle of approximately 5.1° x 3.8°, with a red fixation cross displayed before the image appears. In all tests stimulus presentation was implemented by MATLAB (R2014b, MathWorks) using the Psychophysics Toolbox (58, 59).

#### Time duration discrimination paradigm

To test the effects of prediction-error (PE) on time duration discrimination, a novel 2AFC paradigm was designed in which two images were presented sequentially with 0.5sec delay between them. After the presentation of the second image, participants were asked and had to determine which image had been presented for a longer duration (Fig. 1A). One image was always displayed for a duration of 500 ms – the ‘reference’ duration, whereas the other image was presented for 500ms plus an additional duration (Δt). Because the Just Noticeable time Difference (JND) in time duration discrimination tasks have been reported to range around 15-20% of the standard duration (29, 60), we set Δt in the present experiment to range from 0 ms (equal presentation time for both images) to 133 ms, corresponding to Δt of 0%, 6.6%, 13.3%, 20%, and 26.6% of the 500 ms reference duration.

To generate prediction errors, numbers representing monetary gains and/or losses were presented – overlaid on each image. The first image was always presented with the number zero, while the number of the second image could be negative - generating (PE−); zero - generating no PE (PE0), or positive - generating (PE+). To create a baseline expectation, a large proportion of trials were PE0 trials (60% in the behavior-only group and 80% in the imaging group), with an equally smaller proportion of trials being PE+ and PE− trials. Importantly, the monetary gain or loss did not depend on the performance of the duration discrimination task. For the behavior-only group, PEs were generated by presenting numbers ranging from −5 to +5, in steps of 0.5. For the imaging group, PEs were generated by presenting either −2 (PE−), 0 (PE0), or +4 (PE+). There was a homogenous distribution of presentations across PE values (i.e. all values were presented in a similar number of trials; with the exception of PE0 trials). Moreover, the number of trials for each Δt was equal across PE values, with a counterbalanced order of presentation.

#### Just Noticeable Difference (JND) paradigm

JND estimates for each participant represent the minimal difference in time duration between two stimuli that can still be detected with a high probability. Two psychophysical methods were used to estimate the JND in the present study. First, we used a one-up-two-down staircase procedure (61). Specifically, in each trial two images were presented sequentially (one for 500 ms, the other for 500ms + Δt). If participants correctly discriminated which image was presented for the longer duration, the order of presentation was reversed, and if participants again made a correct discrimination, Δt was decreased by an adaptive amount of time. Whenever an incorrect discrimination was made, Δt was increased by the same amount (up to a maximum of 250 ms). Initial amount was determined on 125 ms, with a 10% decrease of the step size after every trial (regardless of participants’ discrimination accuracy). This process repeats itself until a stopping criterion of correct discrimination after cumulative 6 previous errors have been reached (number of trials was not limited), at which point a threshold has been determined which yields an expected value of 0.707 probability of making a correct discrimination. Of note, while numbers were presented on the images at all times, monetary gains and losses were only associated with the numbers at the test conducted at the end of the experiment (see Procedure below). The second procedure used the method-of-constant-stimuli (MCS) in which a fixed set of pre-determined Δt’s were used (see Time duration discrimination paradigm). Otherwise, the presentation of images was similar as before. The resulting data was then used to generate a psychometric curve for each participant, by fitting a generalized linear regression of the responses to a binomial distribution, and the threshold was defined as the Δt in which participants had a 0.707 probability of making a correct discrimination.

### General procedure

#### Behavior-only group

Following general instructions, a JND time duration discrimination threshold was first estimated for each participant. JND estimate using the staircase procedure was conducted to one image with a zero number, and two additional JNDs were estimated using two MCS procedures of 40 trials each. In one of the MCS procedures, both the first and the second image had the number zero, while in the other MCS procedure the first image was presented with the number zero, while a random number was presented on the second image. This allowed us to control for threshold, visual confounds and value (Fig. S6C). Next, participants performed the time duration discrimination paradigm.

#### Imaging group

The procedure of the imaging group is largely similar to behavior-only group, with the following modification: JNDs were estimated outside the MRI scanner, before and after the main paradigm, and for all images and pairs. In the second estimation of JND (but not the first), after the scan, participants gained and lost money based on the numbers presented on the image, thus allowing us to control for reward value confounds. The time duration discrimination task was performed while undergoing fMRI scanning. Minimum of three training trials were provided inside the scanner to allow participants to become accustomed to the scanner.

### TOE measures from behavior

A well-established perceptual phenomena is the Time Order Error (TOE), which predicts that the duration of the first stimulus in a sequence of stimuli with equal durations, is perceived as longer as compared to succeeding stimuli durations (26, 27).

Time order error can be defined as the difference in probability of successful discrimination as a function of trial type (i.e., Short-Long or Long-Short):

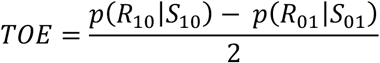

Where *R*_10_ represent a response that the first stimulus is longer, and *S*_10_ represent LS trial, i.e. first stimulus is longer. *R*_01_, *S*_10_ represent the opposite response (i.e. the second stimulus is longer) given a SL trial. To compute the above we extracted individual performance during PE0 trials separately for every Δt, and used the resulted TOEs to correct the probability of discrimination error in all PE-type trials (fig. 1C, Fig. S1D, Fig. S2D, Fig. S3D):

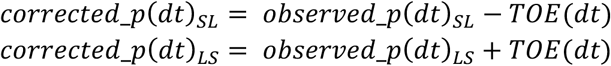

Consequently, positive and negative TOE in time duration discrimination occurs if the first stimulus is perceived as having a longer and shorter duration, respectively. In our experiment, we found positive TOE across all Δt’s (Fig. S4A), thus offering an explanation as to why the probability of making a mistake when Δt = 0 (i.e. the duration of both images is identical) is different than chance level and opposite between LS and SL trials (Fig. S1C, Fig. S2C, Fig. S3C). Specifically, when the duration of the first image was longer (LS trials), TOE causes an even longer perceived duration, generating easier trials (larger perceived Δt). By contrast, when the duration of the first image is shorter (SL trials), TOE causes a shorter perceived duration, leading to more difficult trials (smaller perceived Δt).

### Computational Modeling

Performance (i.e. the probability of making an incorrect discrimination) was modeled as a logistic function where the power of the exponent is linear with respect to Δt (4, 28, 29):

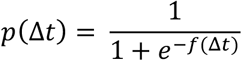

We assume that performance depends linearly on Δt, i.e. larger Δt leads to better performance:

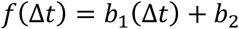

Next, we address other parameters that could influence the perceived Δ*t*.

First, we included the Time Order Error (TOE). As explained above, in our experiment we found positive TOE across all Δt’s (Fig. S4A), which predicts that the duration of the first stimulus is perceived as of longer duration. TOE was here modeled by the parameter *ε*.

Second, the bias in time duration discrimination due to PE, here denoted by the parameter *θ*, is caused by the mismatch between the numbers presented together with the first and the second image, and the expected reward. First, the value of the prediction error on trial *i* was computed as the difference between the actual presented value and current estimation of expected value:

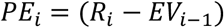

The expected reward (EV) was initialized to 0 and was updated on every trial using a learning-rate parameter α:

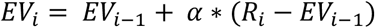

Our results indicate that PE+ and PE− cause the duration of a stimulus to be perceived as longer and shorter, respectively. Accordingly, PE+ decreases performance in LS trials (causing the perceived duration of the second stimulus to be more similar to that of the first stimulus), and vice versa in SL trials for PE−. Incorporating *ε* and *θ* into the model gives the following expression:

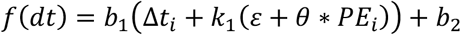

Where

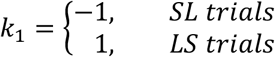

Finally, one additional parameter γ was added to the model to account for a ceiling effect observed in LS trials (in these trials performance is initially much closer to the perceptual threshold (JND) due to TOE, thus might be bounded). The final model looks as follows:

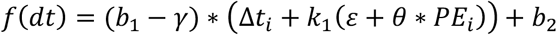

Where

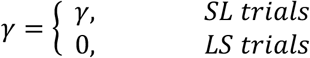

The model is then plugged into the logistic probability function and estimated separately for every subject, as described below.

To estimate the parameters of the model, we computed the maximum a-posteriori (MAP) probability using MATLAB’s function *fmincon* (MathWorks). We assumed uniform prior on the parameters and used bounds as follows:

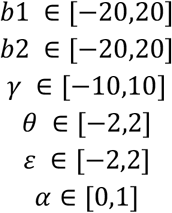

To make sure that results hasn’t been biased due to outliers, we also estimated the MAP with a Gaussian distribution over each parameter with a mean 0 and variance X, where X corresponds to the bound described above for every parameter. Finally, we also computed unconstrained maximum likelihood of the parameters with no priors and no bounds using MATLAB’s function *fminunc*. All results are highly similar with respect to model fit and parameters values.

### fMRI data acquisition

Images were acquired on a 3T Siemens MAGNETOM Tim-Trio scanner. Functional T2* weighted images were acquired using a gradient-echo EPI sequence (TR = 2000 ms, TE = 30 ms, flip angle = 75°, 32 slices with 10% gap scanned in a descending order with phase encoding direction anterior-to-posterior at 30° toward coronal from anterior commissure–posterior commissure (ACPC) plane (63), slice thickness 3 mm, voxel size 3×3×3 mm, FOV 216) in 5 separate scanning sessions (up to two minutes between sessions). Anatomical T1-weighted images were acquired after the functional scans (TR = 2300 ms, TE = 2.98 ms, flip angle = 9°, voxel size 1×1×1 mm, FOV 256). The anatomical scan covered the whole brain while functional scan covered the whole brain except a small area in the dorsal part of the parietal lobe. To improve signal-to noise ratio of the event-related design, order of trials and Inter-Trial-Interval (ITI) in the scanner was determined using OPTSEQ2 (64).

All imaging data were preprocessed and analyzed using Brain Voyager QX 3.4 (Brain Innovation Maastricht, The Netherlands, (65) and MATLAB R2014a (MathWorks) with BVQX/Neuroelf toolbox v1.0 (Jochen Weber, http://neuroelf.net/). Preprocessing included slice scan time correction, motion correction and high-pass filtering. Images were then co-registered and normalized into Talairach space (66) and spatially smoothed with an isotopic 6 mm FWHM Gaussian kernel.

### Statistical analysis

#### Data analysis

Performance on the time duration discrimination task was estimated as the proportion of correct discriminations for each combination of PE, Trial Type, and Δt. These results were then analyzed via repeated measures ANOVAs with these factors and performance as dependent variable. Trials in which no response was made were discarded from analysis.

#### Behavior-only group

First, positive and negative outcomes were grouped into two magnitudes (Fig. S1): small (<|3|) and large (>=|3|). A three-way repeated measures ANOVA was conducted with factors Trial-type (LS, SL), PE type (PE−_large, PE−_small, PE+_large, PE+_small, PE0), and Δt (0-26.6% corresponding to 0-133ms). The dependent variable was task performance, as defined by the proportion of correct time duration discriminations in each condition. We expected and found a significant main effect of Δt (Fig. S1A, Fig. S5, *F*_4,68_ = 26.9, p < 10^−12^). There was no significant interaction between PE type and Δt (*F*_8,136_ = 1.3, p = 0.24), nor was the three-way interaction significant (*F*_8,136_ = 1.18, p = 0.3). Further, there were no significant main effects of PE type (*F*_4,68_ = 0.7, p = 0.59) or of Trial type (SL/LS) (*F*_1,17_ = 4.1, p = 0.06). This means that correct/incorrect time discrimination was not affected by the PE in the specific trials nor by the short-long or long-short structure of it.

Importantly, we found a significant interaction between PE and Trial type (*F*_4,68_ = 9.79, p < 10^−5^; Fig. S1B), in line with our hypothesis and main finding.

Because there was no difference in the result between large and small values (Post-hoc Tukey tests, PE+_small vs. PE+_large; PE−_small vs. PE−_large; all p > 0.12), we grouped values to one positive and one negative group, generating three levels of PE type, i.e. PE+, PE−, PE0 (Fig. S2). Indeed, this did not change the aforementioned findings: no significant main effects of PE type (*F*_2,34_ = 1.5, p = 0.24); no main effect of Trial type (*F*_1,17_ = 4.03, p = 0.06), and replicating the same main finding of significant interaction between PE and Trial type (*F*_2,34_ = 10.89, p = 0.0002; Fig. S2B).

We next validated that the interaction between PE type and Trial type is driven by the same effects as described in the main text. Indeed, we found better performance in SL trials for PE+ vs. PE− (mean PE+ = 0.3, mean PE− = 0.49, post-hoc Tukey, p=0.008) and compared to PE0 (mean PE0 = 0.37, post-hoc Tukey, marginal p = 0.06). Correspondingly, trials with PE− showed worse performance as compared to PE0 (post-hoc Tukey, p = 0.05). In contrast, performance in LS trials during PE+ was worse compared to PE− trials (mean PE+ = 0.39, mean PE− = 0.21, post-hoc Tukey, p = 0.005) and PE0 trials (mean PE0 = 0.275, post-hoc Tukey, p = 0.02). There was no significant difference between PE− and PE0 trials (post-hoc Tukey, p = 0.11). Finally, we normalized to units of individual JND, and found similar results (Fig. S1E,2E), as well as no change in thresholds between different JND measures and the threshold during the main task (Fig. S6).

#### Imaging group

To examine neural correlates of the impact of PE on time duration discrimination, we repeated the experiment with additional subjects (n = 35) while they underwent functional imaging (fMRI). To ensure a stronger PE effect, we increased the proportion of the neutral PE0 trials to 80%, and presented three levels of PE (negative: PE−, neutral: PE0, positive: PE+; remember there was no effect of PE magnitude when comparing 3 to 5 levels). In addition, to avoid loss-aversion effect (Kahneman, 1979) in brain activity, we used twice the magnitude of PE+ compared to PE− (Tom et al., 2007).

Importantly, the main behavioral effect of PE impact on time discrimination was replicated in the scanner as well, and we found a significant interaction between PE and Trial type (*F*_2,68_ = 9.1, p = 0.0003; Fig. S3A, B). In LS trials (mean PE+ = 0.31, mean PE− = 0.23, mean PE0 = 0.28) performance was worse for PE+ as compared to PE− (post-hoc Tukey, p=0.05), and in SL trials (mean PE+ = 0.33, mean PE− = 0.5, mean PE0 = 0.46) performance was better for PE+ as compared to both PE− and PE0 (post-hoc Tukey, p<0.001 for both). There was no difference in performance between PE− and PE0 in SL and between PE+ or PE− and PE0 in LS trials (post-hoc Tukey, p>0.1 for all). These results replicate the behavior in the scanner demonstrating that PE+ and PE− cause stimulus durations to be perceived as longer or shorter, respectively.

Similarly, there was a significant main effect of Δt (*F*_4,136_ = 59.05, p < 10^−5^; Fig. S3A), a significant interaction between Trial type and Δt (*F*_4,136_ = 6.6, p < 10^−5^), and no significant three-way interaction (*F*_8,272_ = 0.8, p =0.59).

#### A behavioral difference between the Imaging and the behavior-only groups and its explanation (or: why does PE0 also have a slight bias in the fMRI group)

There was one difference between the imaging group and the behavior-only group: in the Imaging group we found significant main effects of PE type (F_2,68_ = 8.2, p = 0.0006) and Trial type (F_1,34_ = 24.2, p < 10^−4^), as well as a significant interaction between PE type and Δt (F_8,272_ = 2.4, p = 0.01). The main effect of PE stems from better performance in PE+ compared to PE− (PE− vs. PE+; post-hoc Tukey, p=0.02) and PE0 (PE+ vs - PE0; post-hoc Tukey, p=0.001).

We hypothesized that a possible explanation can be related to the differences in the outcomes assigned to each group. First, both the values (−2 and +4) and the range (6) in the imaging group outcomes were smaller as compared to behavior-only group (extremes: −5 to +5, range: 10). This reduction in the magnitude of PE+ and PE− could have attenuated the impact of PE+ and PE− on discrimination performance, as compared to PE0. Second, even though 80% of trials were PE0 trials, the expected outcome could have been biased by the magnitude difference between PE+ and PE− (i.e. the expected value across the study is actually +0.2, compared to zero-0 in the behavior-only group). Accordingly, the PE0 would actually represent a slightly negative PE for the imaging group (as opposed to the behavior-only group), causing PE0 trials to be perceived as of negative value, at least in part (Fig. S4, see comparison between groups below). This, in turn, could indicate why performance in those trials is similar and not significantly different than PE− trials (PE− vs-PE0; post-hoc Tukey, p=0.98).

Interaction between PE type and Δt results from better performance in PE+ compared to PE0 trials in “easy” trials (Fig.S5, Δt levels 3-5, corresponds to Δt >= 13.3%) but not “hard” trials (Δt levels 1-2, corresponds to Δt < 13.3%) (Δt3; PE+ vs. PE0, post-hoc Tukey, p=0.04; Δt4; PE+ vs. PE0, post-hoc Tukey, p=0.002; Δt5; PE+ vs. PE0, post-hoc Tukey, p=0.004). All other pair-wise comparisons weren’t significant except one (Δt4; PE+ vs. PE−, post-hoc Tukey, p=0.01), indicating that PE0 trials are more similar to PE− trials as Δt increases, strengthening the explanation above and in main text.

In addition, main effect of trial type was found, a result of better performance in LS vs. SL trials, likely stemming from same reasons explained above. There was also a higher probability for mistake during PE0 trials for the imaging group (Fig. S4C, two-samples t-test; *t*_51_ = 2.52, p = 0.01), further confirming the difference in expected value for PE0 between the groups (i.e. PE0 was likely slightly negative in the Imaging group).

Importantly, when comparing the two groups (with and w/o imaging), we did not find any difference in performance for the PE+ and PE− in both SL or LS trials (*t*_51_ = 0.8, p = 0.4; *t*_51_ = 0.9, p = 0.37). Finally, we normalized to units of individual JND as before, and found similar main results (Fig. S3E), as well as no change in thresholds as a result of the discrimination task (Fig. S6).

#### JND measures

To control for changes in individual thresholds we compared JND obtained using staircase method, JND extracted from method-of-constant-stimuli (MCS) curve generated before main paradigm (see methods) and JND extracted from main paradigm PE0 curve, all computed to the same criteria (proportion of discrimination errors). For the behavior-only group, results indicate no difference between JNDs (*F*_2,46_ = 0.7, p = 0.5) which validates the use of staircase JND to normalize the results (fig. S6), indicating that the bias found in the perception of time was not confounded by differences in perceptual threshold between trials or a change in threshold during the paradigm.

For the imaging group, analyzing JND also indicates no difference in time discrimination threshold within different images at the beginning or at the end of the experiment (one-way ANOVA; JND at the beginning *F*_4,120_ = 0.6, p = 0.66, JND at the end *F*_4,120_ = 0.3, p = 0.8), and no change in JND from beginning to end of experiment was found (paired t-tests, p > 0.09, Fig. S6).

Finally, to make sure that indeed individual thresholds did not confound the results, we repeated the experiment with another 12 participants and measured JND before and after main paradigm. Threshold (JND) of different images and trials was found to be similar (before main paradigm *F*_4,55_ = 0.5, p = 0.7, after main paradigm *F*_4,55_ = 0.27, p = 0.89) and did not change due to main paradigm (paired t-tests, p > 0.1, fig. S6)

#### fMRI data analysis

Analyses consist of random effects Analysis of variance (ANOVA) based on general linear models (GLM), with event regressors defined from the onset of the first image until the offset of the second image, models as box-car functions and convolved with a canonical hemodynamic response function (HRF). All models included 6 regressors to account for head movements and a regressor to account for the motor response, modeled from the onset of the cue until the subject’s response.

Multiple comparison correction on cluster size was done using non-parametric permutation test (30, 31). Null distribution of maximal cluster size was built separately using 1000 iterations for every analysis (PE x Correctness interaction, all trials with probability weights as parametric modulators). On every iteration, all labels of all trials were randomly shuffled for every subject, and a GLM for every voxel was computed using the same definitions as in main analysis. Finally, ANOVA model was created and maximum cluster size was extracted (direct or diagonal proximity in one dimension was sufficient to include voxels in same cluster) using the MATLAB function *bwlabeln*. Cluster Defining Threshold (CDT) level was set on p < 0.005. Analysis using trial-to-trial PE estimates as parametric modulations consist of separate contrasts for PE types and corrected for multiple comparisons using false discovery rate of q < 0.05.

#### Comparison between different parts of the experiment

To test the intention and interpretation that PE0 trials are biased toward the expected value in the study, we analyzed both halves of the trials separately within each group. First, we made sure that equal number of trials was used for every half. Second, we fitted the corresponding proportion of discrimination errors of every subject to a logit function corresponds to a psychometric curve, to be able to analyze the different slopes of the function. If indeed PE0 trials are biased towards the expected value (e.g., considered a small negative PE in the imaging group, while remain PE0 in behavior-only group), we expected to see that the slope of the psychometric curve would shift as participants gather more evidence to compute the expected value. This means that the second half of trials in the imaging group are expected to have higher proportion of discrimination errors as a function of Δt compared to the first half, while no other difference is expected. Moreover, we expected no significant differences between first and second half of trials in all trial-types.

As hypothesized, we found a significant decrease in psychometric curve of second-half PE0 trials of the imaging group compared to first half (fig. S4C; paired t-test; *t*_34_ = −2.5, p = 0.01) and no change in performance for PE+ (*t*_34_ = 0.01, p = 0.98) or PE− trials (*t*_34_ = 0.2, p = 0.8). In behavior-only group, no difference in performance between first and second half of the study was found (PE0; *t*_17_ = 0.3, p = 0.7. PE+; *t*_17_ = 0.47, p = 0.6. PE−; *t*_17_ = 1.4, p = 0.15). Taking together these results support the notion that participants in the imaging group may have considered PE0 trials to be negative PE as the number of trials increased.

#### Computational model selection

Our computational models were used to track of trial-to-trial probability of discrimination error, corresponds to the behavioral measure used in analyses. The selected model (here model 1) described in main text is a follow:

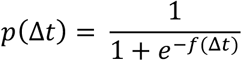

With

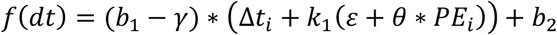

We tested 5 variations of the model:

Model 1 – as described, total of 6 parameters (*b*_1_, *b*_2_, *α, ε, θ, γ*)
Model 2 – no free parameter *b*_2_ – total of 5 parameters
Model 3 – no *γ* to correct for ceiling effects – total of 5 parameters
Model 4 – no free parameter *b*_2_, no *γ* - total of 4 parameters
Model 5 – as model 1, with separate parameters for the bias due to PE+ (*θ*+) and PE (*θ*-) - total of 7 parameters

For each model, mean AIC (Akaike Information Criterion) was estimated using the computed likelihood of the model for each subject regulated by the number of parameters. Mean AICs were used as model evidence in a Bayesian Model Selection for group studies (BMS) (62), in which the exceedance probabilities of all models (i.e., how likely each model is to explain the obtained model evidence) is estimated. Results indicated a 0.98 probability for model 1 to be the model best explaining the evidence, therefore it is the selected model.

#### Assigning PE type to trials according to selected model

In order of generating different trial types according to the modeled PE, trials were categorized based on the estimated PE. Since almost no trials estimated precisely PE = 0, and in order of assigning enough trials for all PE types to generate a valid statistic, we selected a threshold for which a trial with absolute value of PE smaller than that threshold would be assigned as PE0 trial, and PE larger or smaller than the signed threshold would be assigned as PE+ and PE− trials, respectively:

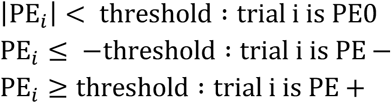

A threshold of 0.1 was chosen since it produced a similar proportion of PE0 trials as in the actual design. To make sure that this selection did not bias results, we also tested thresholds of 0.2, 0.5, 0.01, and 0.005. All tests resulted in same main effects and interactions except the three-way interaction that wasn’t significant for low (below 0.05) thresholds. This indicates a robust PE type statistic across different levels of classification.

#### Correlation between estimated and observed TOE

Our computational model assumes one measure of TOE from both trial types and across Δt. To test correlation between measured TOE and estimated TOE from our model, TOE from PE0 curve was measured by finding the bias from 0.5 probability for discrimination error in trials of Δt = 0, separately for LS and SL trials. We then computed the average TOE from curve, i.e. the average across trial-type, and correlate that with the estimated TOE (fig. 2C). To make sure that the correlation is stable we computed the correlation for each trial type separately and found that it remained significant in behavior-only group (Pearson correlation coefficient; SL trials; r = 0.8, p = 0.0008. LS trials; r = −0.71, p = 0.005) and imaging group (SL trials; r = 0.66, p = 0.0004. LS trials; r = −0.69, p = 0.0002). Notice that the correlation is opposite for SL and LS trials, since TOE biases performance in opposite directions.

#### Correlation between individual interaction activations and behavioral bias

We calculated an interaction-score for each subject’s Putamen activity and correlated it with the individual behavioral bias (θ) derived from the model:

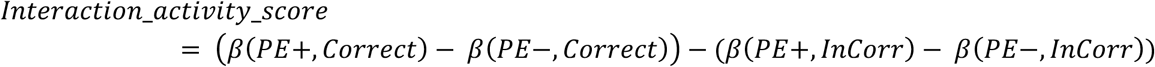

Where *β*(*PE+, Correct*) is the average regional activity in trials of PE+ with correct response, and so forth. The interaction_activity_score provides an approximate individual quantification of the interaction found in the imaging analysis (PE_TYPE * performance).

## Tables

**Table S1.**
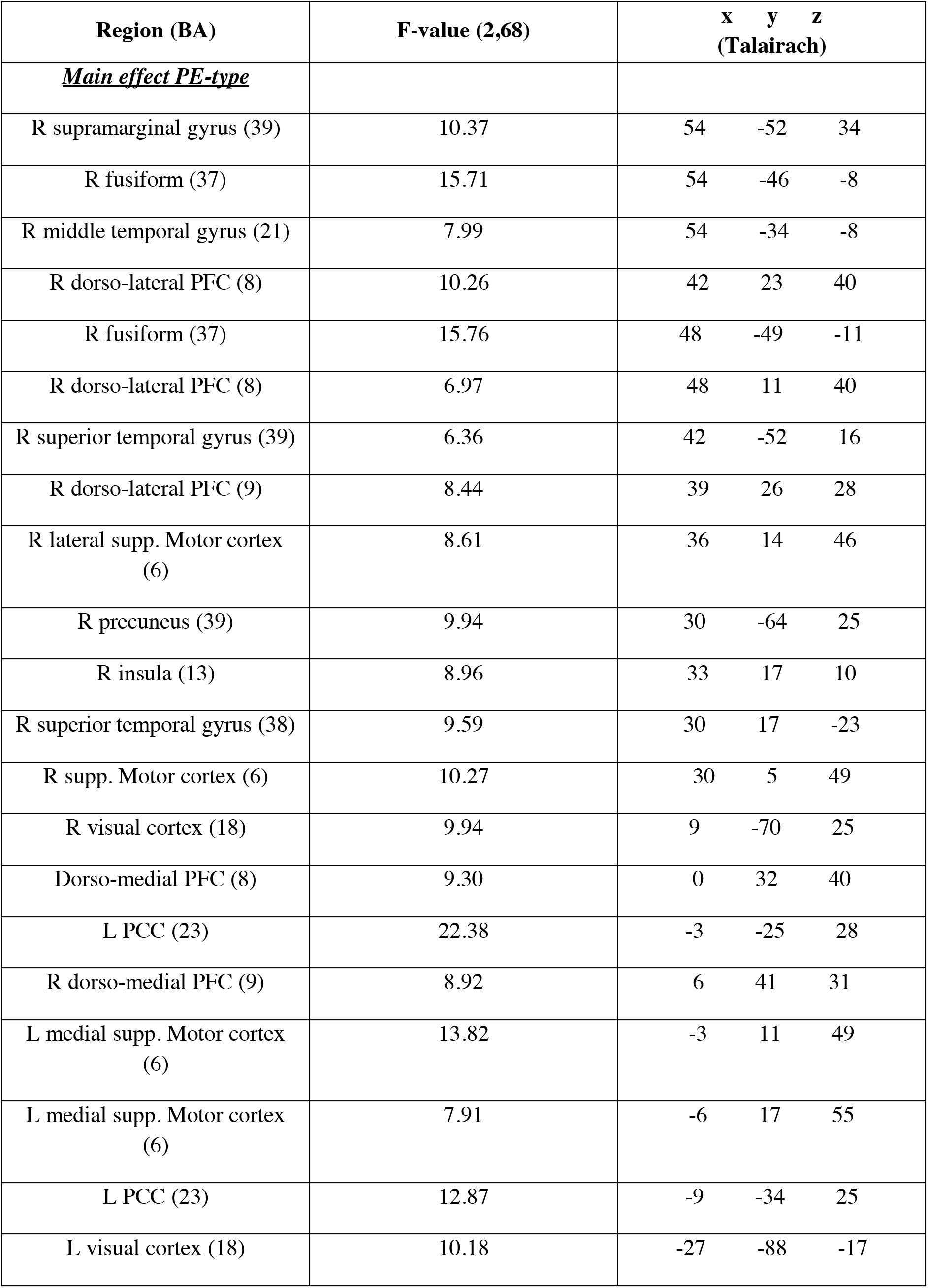
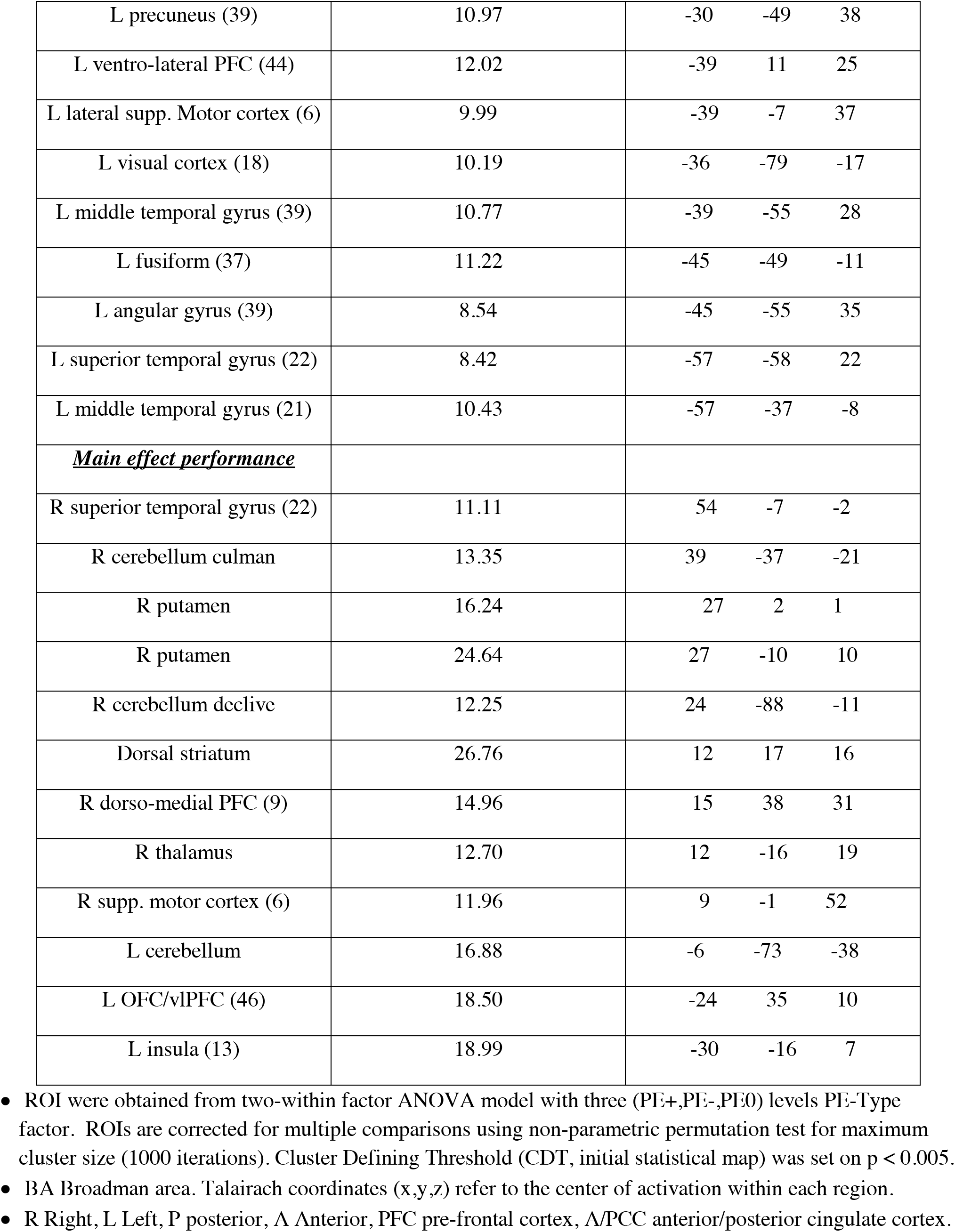
Main effects of PE-type x Performance.

**Table S2.**
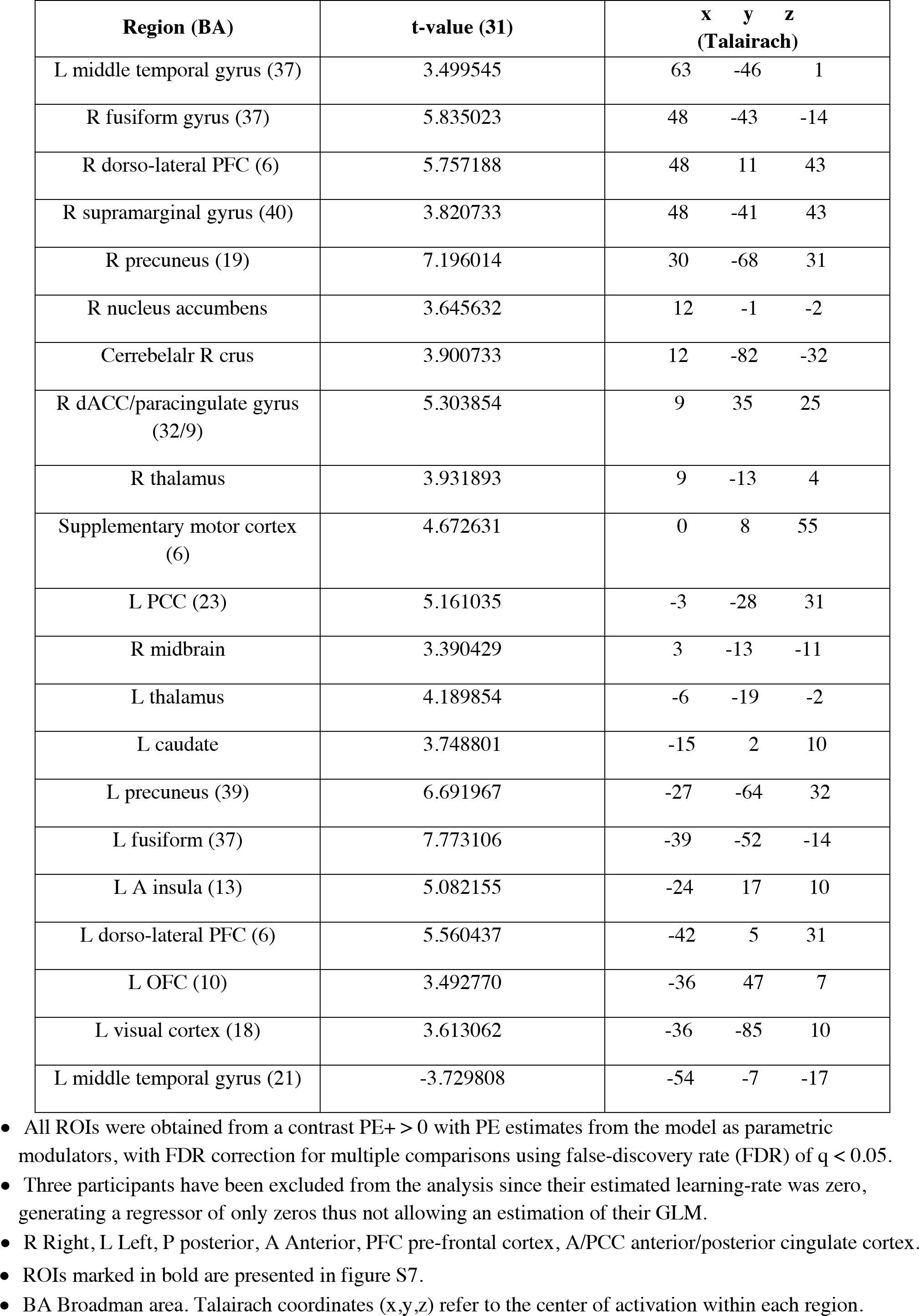
Activation correlated with trial-to-trial PE estimation.

**Table S3.** Activations correlated with trial-to-trial probability of correct time discrimination.

| **Region (BA)** | **t-value (34)** | **x y z**<br>**(Talairach)** |
| --- | --- | --- |
| R inferior frontal gyrus (6) | 4.41 | 46 3 32 |
| **R A ventro-lateral PFC (47) /**<br>**OFC** | -4.5 | 26 27 -11 |
| **R ventral ACC (25) / Nucleus**<br>**accumbens** | -3.9 | 9 10 -7 |
| **L ventral ACC (25) / Nucleus**<br>**accumbens** | -3.7 | -8 13 -14 |
| L caudate tail | 3.8 | -16 -35 21 |
| **L A ventro-lateral PFC (47) /**<br>**OFC** | -4.1 | -42 29 -6 |

- ROIs were obtained using trial-to-trial estimated probability for discrimination error and corrected for multiple comparisons using non-parametric permutation test for maximum cluster size (1000 iterations). Cluster Defining Threshold (CDT, initial statistical map) was set on $p < 0.005$ . - Talairach coordinates (x,y,z) refer to the center of activation within each region. - ROIs in bold are displayed on figure 3. - R Right, L Left, P posterior, A Anterior, PFC pre-frontal cortex, A/PCC anterior/posterior cingulate cortex.

**Figure S1.**
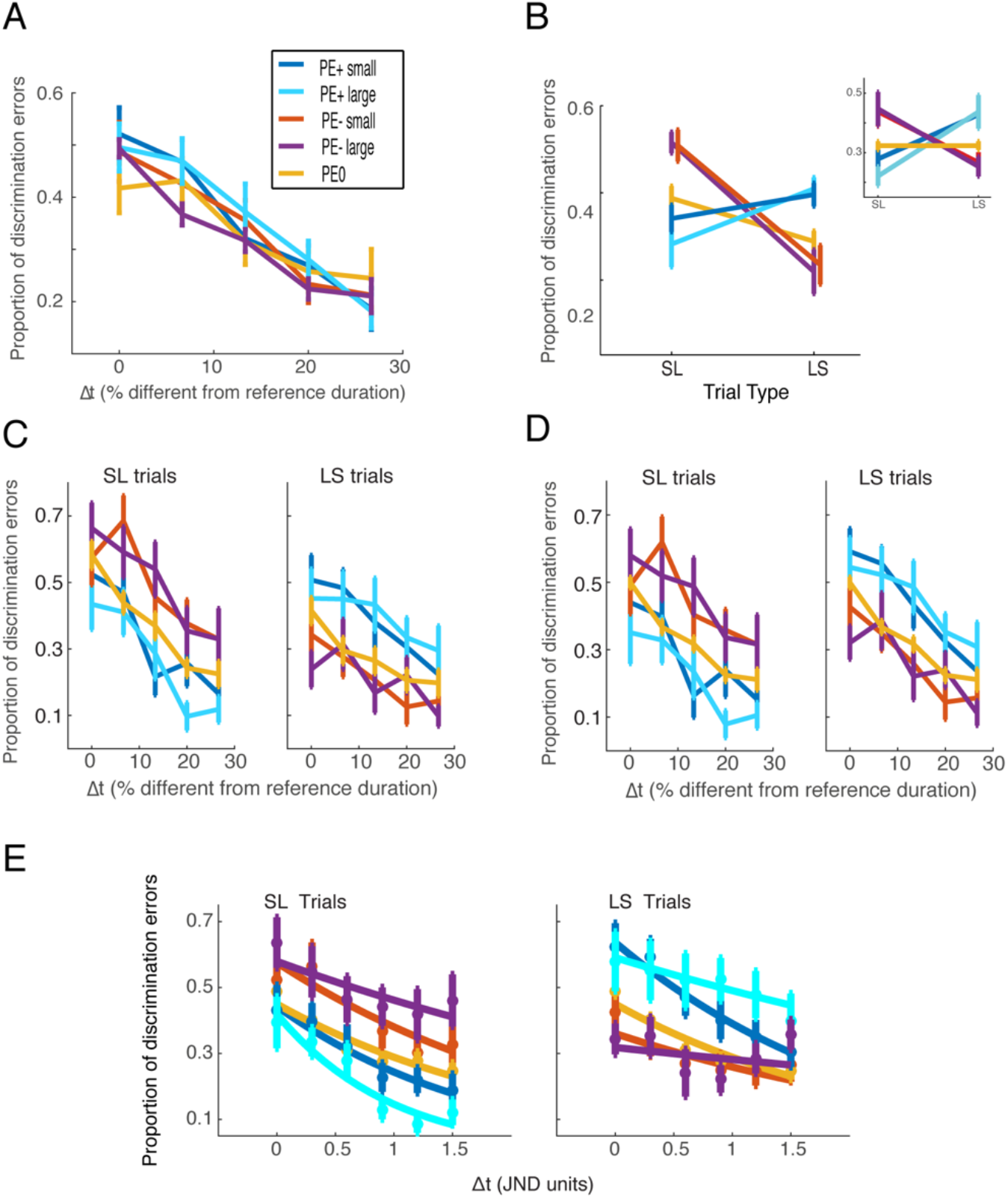
Behavior-only group analyses of performance, separated into 5 levels of PE. Data is presented similar to main Fig.1, but only for subjects in the behavior-only group, when separating trials into 5 levels of PE. Data represented as mean ± SEM. See Supp.Info for Stats. (A) Proportion of discrimination errors as a function of PE type and time difference between images (Δt) in all trials, indicating no difference between PE types. (B) Proportion of discrimination errors as a function of PE type and trial type. An interaction between PE-type and trial-type is evident as PE+ and PE− bias performance in opposite directions relative to PE0 trials. Inset: after correction for individual TOE. (C) Proportion of discrimination errors as a function of time difference between images (Δt), presented separately for Long-short and Short-long trials and corrected for Time-Order Error (TOE). PE+ and PE− bias performance in opposite directions relative to PE0, for all values of Δt. Different outcome magnitudes are not significantly different. (D) Proportion of discrimination errors as a function of time difference between images (Δt), presented separately for Long-short and Short-long trials and corrected for Time-Order Error (TOE). PE+ and PE− bias performance in opposite directions relative to PE0, for all values of Δt. (E) Proportion of discrimination errors as a function of subjective JND, corrected for TOE. Data fitted to a logistic function after JND normalization

**Figure S2.**
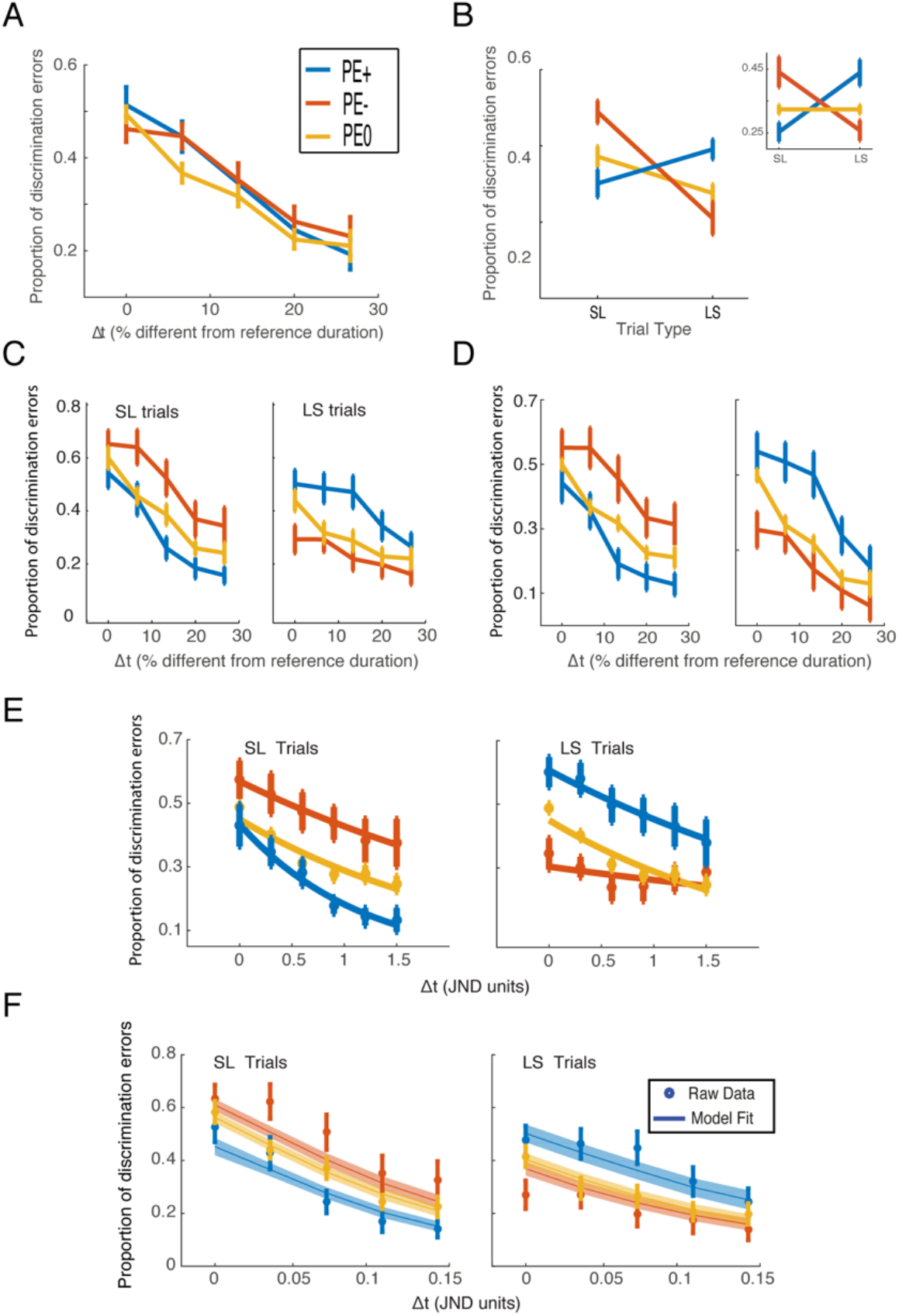
Behavior-only group analyses of performance, separated into 3 levels of PE. Data is presented similar to main Fig. 1 and Fig.S1, but only for subjects in the behavior-only group, when separating trials into 3 levels of PE. Data represented as mean ± SEM. See Supp.Info for Stats. (A) Proportion of discrimination errors as a function of PE type and time difference between images (Δt) in all trials, indicating no difference between PE types. (B) Proportion of discrimination errors as a function of PE type and trial type. An interaction between PE-type and trial-type is evident as PE+ and PE− bias performance in opposite directions relative to PE0 trials. Inset: after correction for individual TOE. (C) Proportion of discrimination errors as a function of time difference between images (Δt), presented separately for Long-short and Short-long trials and corrected for Time-Order Error (TOE). PE+ and PE− bias performance in opposite directions relative to PE0, for all values of Δt. Different outcome magnitudes are not significantly different. (D) Proportion of discrimination errors as a function of time difference between images (Δt), presented separately for Long-short and Short-long trials and corrected for Time-Order Error (TOE). PE+ and PE− bias performance in opposite directions relative to PE0, for all values of Δt. (E) Proportion of discrimination errors as a function of subjective JND, corrected for TOE. Data fitted to a logistic function after JND normalization (F) Model fit to individual behavioral data, averaged over subjects.

**Figure S3.**
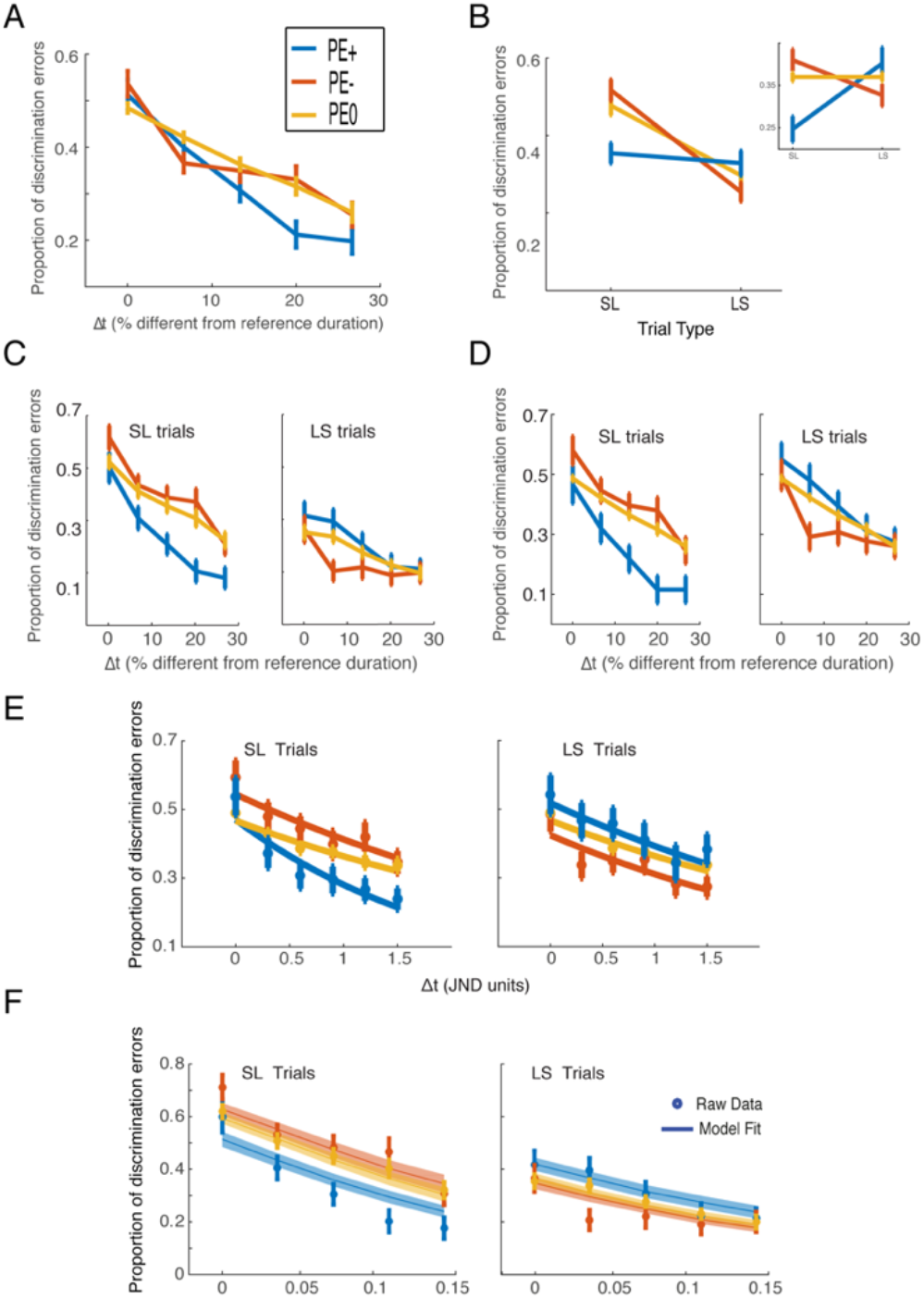
Imaging-group analyses of performance. Data is presented similar to main Fig. 1 and Fig.S1,S2, but only for subjects in the Imaging (fMRI) group. Data represented as mean ± SEM. See Supp.Info for Stats. (A) Proportion of discrimination errors as a function of PE type and time difference between images (Δt) in all trials, indicating no difference between PE types. (B) Proportion of discrimination errors as a function of PE type and trial type. An interaction between PE-type and trial-type is evident as PE+ and PE− bias performance in opposite directions relative to PE0 trials. Inset: after correction for individual TOE. (C) Proportion of discrimination errors as a function of time difference between images (Δt), presented separately for Long-short and Short-long trials and corrected for Time-Order Error (TOE). PE+ and PE− bias performance in opposite directions relative to PE0, for all values of Δt. Different outcome magnitudes are not significantly different. (D) Proportion of discrimination errors as a function of time difference between images (Δt), presented separately for Long-short and Short-long trials and corrected for Time-Order Error (TOE). PE+ and PE− bias performance in opposite directions relative to PE0, for all values of Δt. (E) Proportion of discrimination errors as a function of subjective JND, corrected for TOE. Data fitted to a logistic function after JND normalization (F) Model fit to individual behavioral data, averaged over subjects. Notice that in this group (fMRI), it seems that PE0 also has a slight bias, because the distance between PE+ and PE0 is not similar to the distance between PE− and PE0 (can be seen in B,C,D,E). This can be explained by the different values used in this experiment (−2,4), that create an expected value of 0.2 (on average), and hence PE0 also has a small negative PE. For a full explanation and direct experimental evidence, see subsection “A behavioral difference between the Imaging and the behavior-only groups and its explanation (or: why does PE0 also have a slight bias in the fMRI group)”, and the references there to Fig.S4,S5.

**Figure S4.**
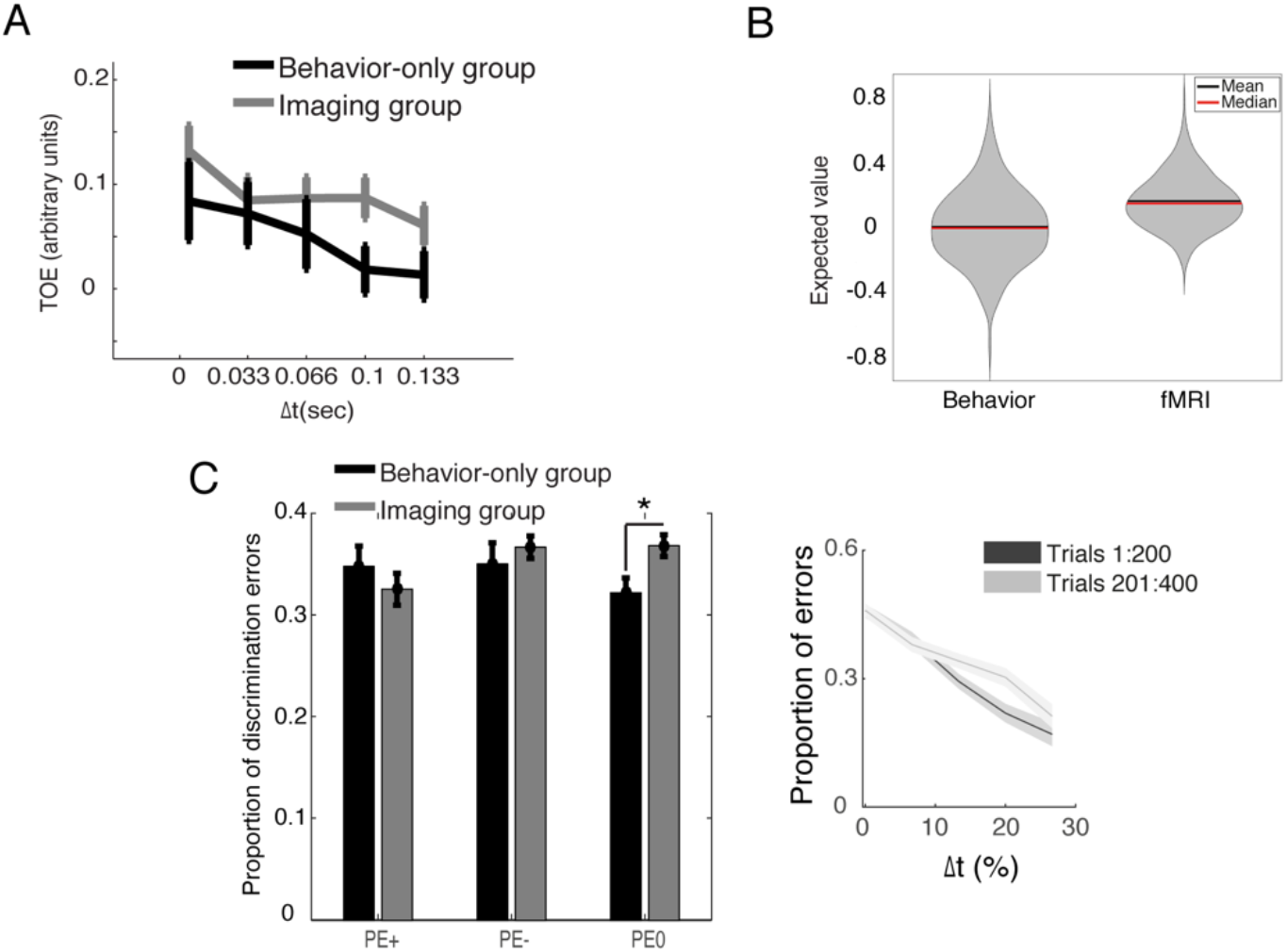
Comparing behavior-only and imaging group. (A) Time order error (TOE) as a function of Δt, measured directly from the behavior. TOE is positive in both groups across all Δt, indicating a positive bias, namely the first stimulus is perceived as longer. (B) Mean expected value from the model in the last 50 trials. Overall, a higher expected value can be seen in the imaging group, likely due to the difference in magnitude between the positive and negative outcomes (4/-2) used for that group. (C) Proportion of discrimination errors grouped by the estimated PE from the model. No difference in performance is seen between the groups for PE+/-. However, higher proportion of errors in PE0 was found in the imaging group, suggesting a non-zero value assigned to these trials during the main paradigm likely due to the difference in magnitude between the positive and negative outcomes (4/-2) used for that group. Right: proportion of discrimination errors for PE0 trials in first and second half of main paradigm in the imaging group. A change in slope with a more moderate slope (namely worse discrimination) for later trials, indicating again that the expected value develops to be non-zero because of the unbalanced negative and positive outcomes.

**Figure S5.**
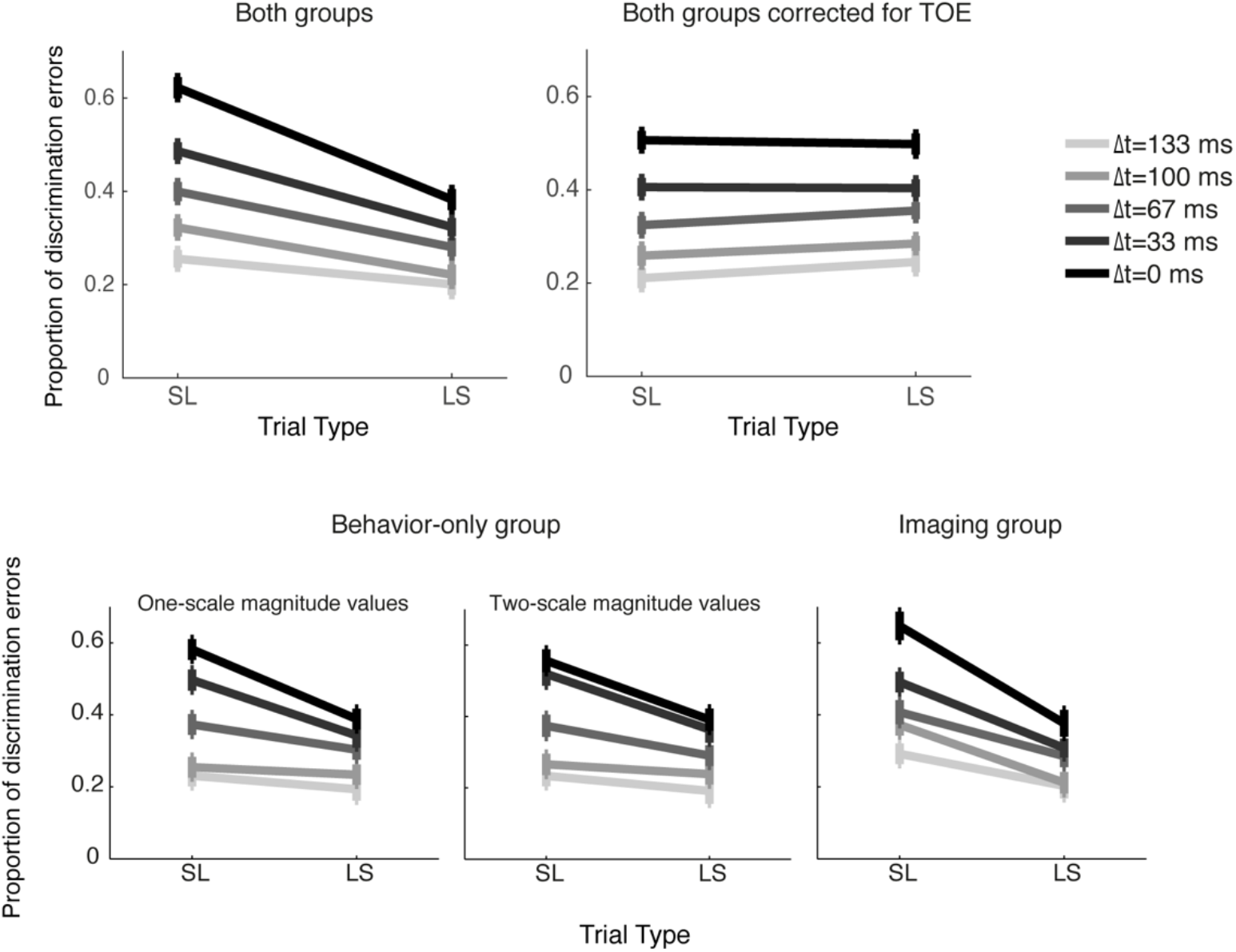
Proportion of discrimination errors and TOE correction. Presented as a function of trial type (Long-Short or Short-Long) and (Δt), demonstrating higher difference in performance between LS and SL trials for small Δt (“hard” trials) compared with higher Δt (“easy” trials). Data is represented as mean ± SEM. Top – all participants (left), data corrected for TOE (right). Bottom – separated according to group (before correction to TOE). TOE occurred in all groups, and was therefore corrected for all (see top-right).

**Figure S6.**
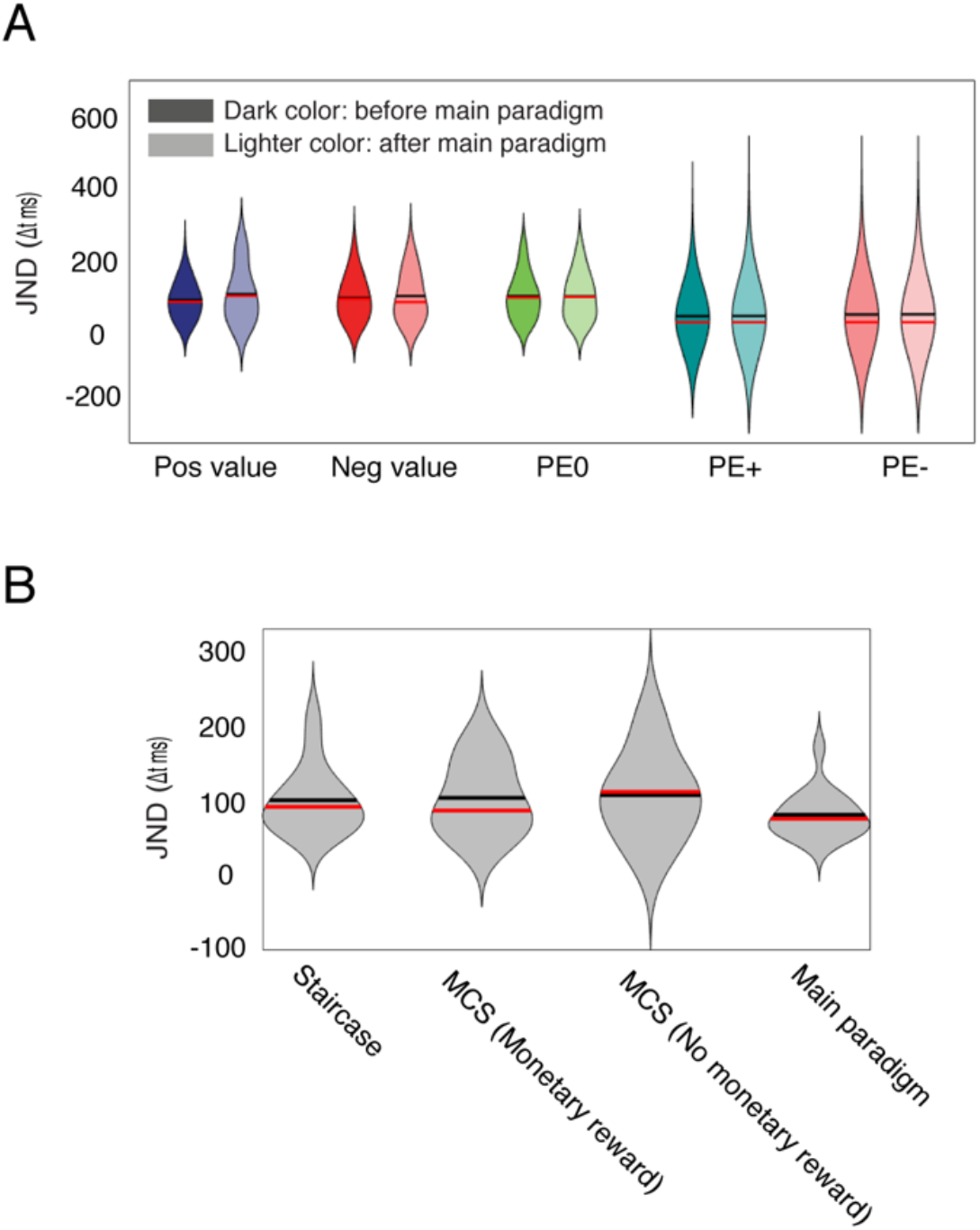
Stability of perceptual thresholds and value control. Data is represented as mean ± SEM. (A) Comparison between JND before (left darker bars) and after (right lighter-color bars). The two sets of left bars measure JND when the two consecutive images were overlaid with the same number (positive or negative), to control for value and visual differences. The middle (PE0) and right set of bars (PE+/PE−) use two images identical to the trial types examined in the main paradigm. Before the paradigm numbers did not have any meaning (i.e. did not provide reward/loss), while at the end they did (like in the main paradigm). Overall, the different conditions show that perceptual thresholds of did not change: before to after the main paradigm, with or w/o real outcome, and with outcome that is predicted (i.e. no PE). These controls suggest that what we observed during the main paradigm is due to PE. (B) Comparison of four different methods for computing individual JNDs, indicating no difference between the approaches (see methods for the different approaches).

**Figure S7.**
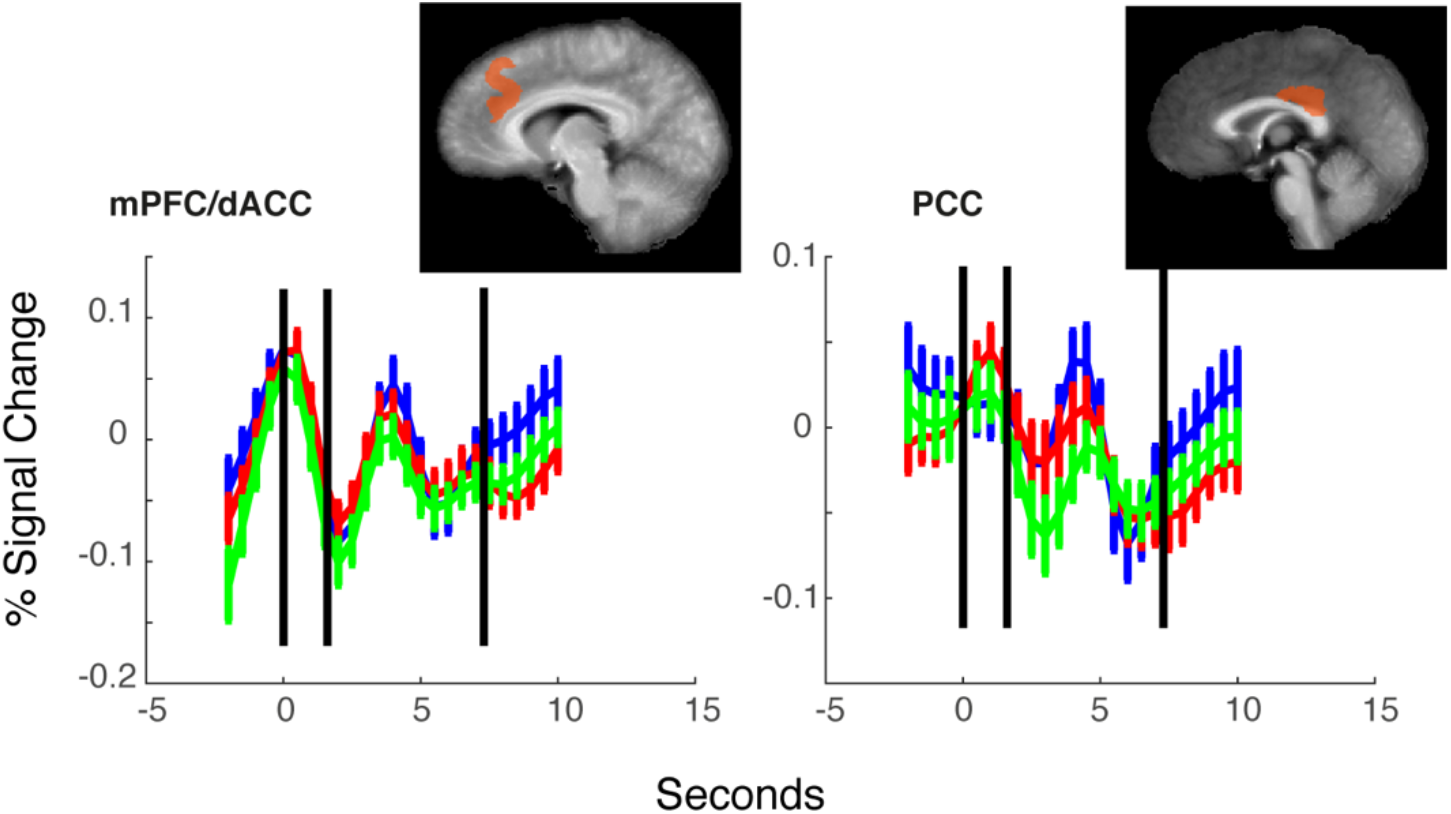
Additional ROIs for trial-to-trial prediction error. Shown are activations in the dACC and PCC. Displayed on average brain. Time courses represent mean % signal change extracted from the ROIs ± SEM. Black vertical lines represent trial onset, trial offset and average onset of next trial, respectively. Activation was set to statistical threshold of q < 0.055 for visualization.

**Figure S8.**
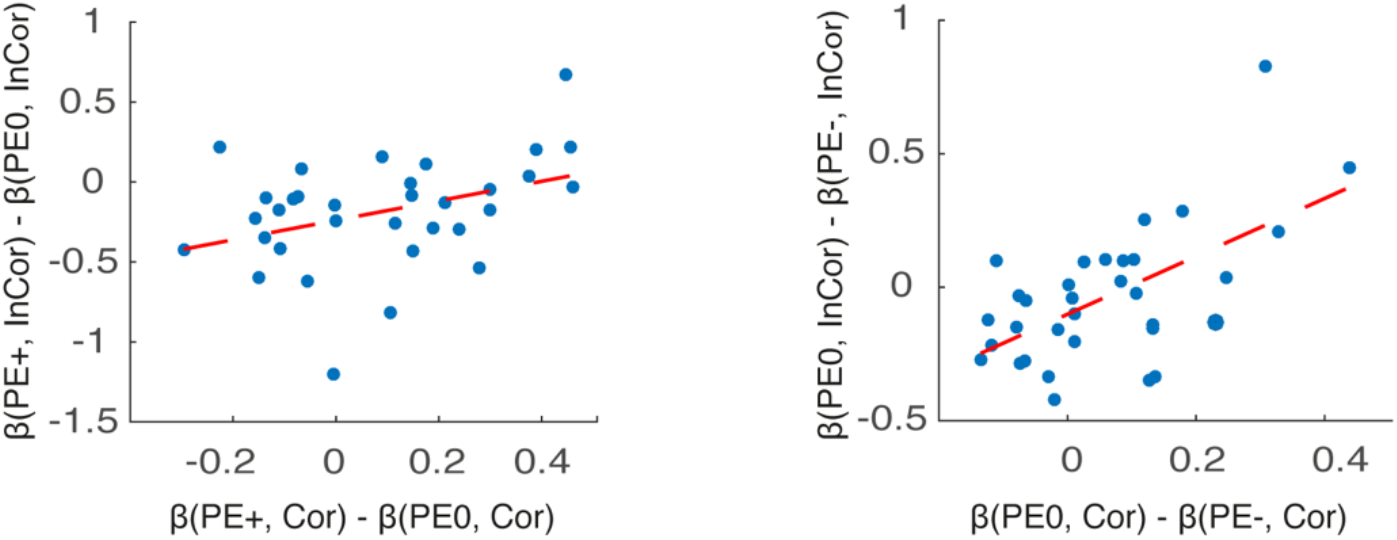
Individual dACC activations for PE+/PE−/PE0 in incorrect vs. correct trials. Individual subject dACC activations (extracted from the ROIs in Fig.3B) for PE+/PE0 in incorrect vs. correct discrimination trials (left), and for PE0/PE− in incorrect vs. correct discrimination (right). Both were significantly correlated, indicating that dACC activity tracks these factors at an individual level as well.

